# Autologous humanized mouse models of iPSC-derived tumors allow for the evaluation and modulation of cancer-immune cell interactions

**DOI:** 10.1101/2021.04.02.437733

**Authors:** Gaël Moquin-Beaudry, Basma Benabdallah, Damien Maggiorani, Oanh Le, Yuanyi Li, Chloé Colas, Claudia Raggi, Benjamin Ellezam, Marie-Agnès M’Callum, Dorothée Dal Soglio, Jean V. Guimond, Massimiliano Paganelli, Elie Haddad, Christian Beauséjour

## Abstract

Modeling the tumor-immune cell interactions in humanized mice is complex and limits drug development. Here, we generated easily accessible tumor models by transforming either primary skin fibroblasts or iPSC-derived cell lines injected in immune-deficient mice reconstituted with human autologous immune cells. Our results showed that either fibroblastic, hepatic or neural tumors were all efficiently infiltrated and partially or totally rejected by autologous immune cells in humanized mice. Characterization of tumor immune infiltrates revealed high expression levels of the dysfunction markers Tim3 and PD-1 in T cells and an enrichment in regulatory T cell suggesting rapid establishment of an immunosuppressive microenvironment. Inhibition of PD-1 by Nivolumab in humanized mice resulted in an increased immune cell infiltration and a slight decrease in tumor growth. We expect these versatile and accessible cancer models will facilitate preclinical studies and the evaluation of autologous cancer immunotherapies across a range of different tumor cell types.

**SIGNIFICANCE STATEMENT:** Preclinical models capable of providing an environment where human tumors are confronted with autologous immune cells are not easily accessible and limit drug development. As an alternative we generated genetically-defined tumor cell lines from primary and iPSC-derived cells for the evaluation of cancer-immune cell interactions in autologous humanized mice.

## INTRODUCTION

Cancer drugs have the worst likelihood of approval when compared to the rest of the industry (1). This situation underlines the need for better preclinical models to emulate clinical conditions in a reliable manner (2). This is particularly true in the era of immuno-oncology, where cancer is no longer viewed as a cell-autonomous disease. Instead, the whole tumor microenvironment, especially the host’s immune system, is now considered as an important modulator of cancer development and elimination through cancer immunoediting mechanisms (3,4). While classical cancer therapies have been shown to indirectly promote anticancer immune response (5,6), new immunotherapies such as checkpoint blockade inhibitors, vaccination or chimeric antigen receptor (CAR) cell therapy aim at enhancing, potentializing or inducing the host’s anti-tumor immune response. However, very few preclinical models manage to provide a relevant immunological environment where human tumors are confronted with autologous immune cells.

Genetically engineered mouse models of cancer are valuable platforms but their translational potential is limited by genetic and physiological differences between mice and humans (7,8) and historically have demonstrated poor translational robustness (9). This led to the development of various chimeric mouse models that use immunodeficient mice NOD/SCID/IL2Rγ^null^ (NSG) as vessels for human tumor growth. While patient-derived xenograft (PDX) models are very potent tools to study established immune system-evading tumors, their use as preclinical platform has some caveats such as maintenance, genetic drift and most importantly their difficult combination with an autologous immune system. Some recent studies have tackled this issue by combining PDX with patient-derived hematopoietic stem cells (HSCs) or reinfusion of in vitro-expanded tumor-infiltrating lymphocytes (10,11). However, the reliance on patient-derived immune cells limits the accessibility and scalability of such models. Similarly, while cancer cell lines are easy to use, they are debatably reproducible (12,13) and suffer the same complexities when it comes to studying autologous tumor-immune system. To address this, HLA matching has been successfully attempted but remains technically challenging (14).

Multiple approaches of mouse humanization are currently being used (15). In humanized adoptive transfer (Hu-AT) models, mature and functional peripheral blood mononucleated cells (PBMCs) from donors are injected to immunodeficient mice for rapid and efficient reconstitution, albeit at the cost of a rapid graft versus host disease (GvHD) onset (16). Alternatively, humanization models using HSCs provide a long term robust human lymphocyte reconstitution (17,18). However, effector T cells are trained on murine thymic tissue, undermining their ability to mount specific autologous TCR interactions. To address this, humanized bone marrow/liver/thymus models (Hu-BLT) employ fetal liver-derived HSCs with surgical implantation of autologous thymic tissue under the renal capsule for improved T cell education (19,20).

Here we propose an approach combining the flexibility of cancer cell lines obtained by transforming primary fibroblasts or iPSC-derived cells with a set of defined oncogenes with the easy access to autologous immune cells from healthy donors. Using either Hu-BLT or Hu-AT mice, we have developed versatile and accessible preclinical models that are uniquely positioned to study immune-naive tumors in an autologous immune setting.

## RESULTS

### Engineered fibroblastic tumors are recognized by autologous immune cells in Hu-AT and Hu-BLT mice

In order to establish tumor models with an easy access to autologous immune cells, we first elected to generate tumor cell lines from skin fibroblasts derived from healthy adult donors. Transformation of fibroblasts was achieved by successive lentiviral transductions of hTERT, SV40ER and HRas^v12^ genes which were shown to efficiently transform human cells (21). These tumor cells were also tagged with the mPlum fluorescent marker for *in vivo* imaging and hereinafter designated 4T cell line. The subcutaneous injection of tumor cells in the flank of NSG-SGM3 mice lead to the formation of tumors in all mice within 3-4 weeks (Fig. 1A). Previous work from our laboratory showed that adoptive transfer of PBMCs in NSG-SGM3 is highly effective at rejecting tumor and allogenic myoblasts (22,23). These naïve tumors were infiltrated by immune cells and partially or fully rejected following the adoptive transfer of 5×10^6^ autologous human (Auto-AT) PBMCs and granulocytes (Fig 1A, B). Previous work from our laboratory showed that injecting more PBMCs does not enhance tumor clearance (23). Flow cytometry analysis of the immune infiltrate of residual tumors showed multiple immune cell populations, mostly restricted to T and NK cells (Fig. 1C and S1). Differential clustering analysis between human circulating and tumor-infiltrating immune cells (hTIIC) suggests that the tumor microenvironment can significantly alter the immune phenotype (Fig. 1C right). For example, tumors were enriched for CD14^+^cells and immunosuppressive CD4^+^CD25^+^CD127^−^Treg cells while depleted of effector CD56^+^ cells, CD8^+^ T cells and CD45RO+ effector memory T cells (Fig. 1D). However, no significant change in the proportion of dysfunctional T cells (defined as positive for the exhaustion markers Tim3 and PD-1) was found, with all mice showing high levels of dysfunctional cells in both blood and the tumor environment (Fig. 1E). Still, we observed variations in the expression level (as detected by mean fluorescent intensity) of the exhaustion markers Tim3 and PD-1 on T cells in both blood and tumors (Fig. 1F). These results suggest that while immune-naïve tumors are efficiently recognized by autologous immune cells, tumors are quickly able to induce an immunosuppressive microenvironment and avoid complete elimination in most cases.

**Figure 1.**
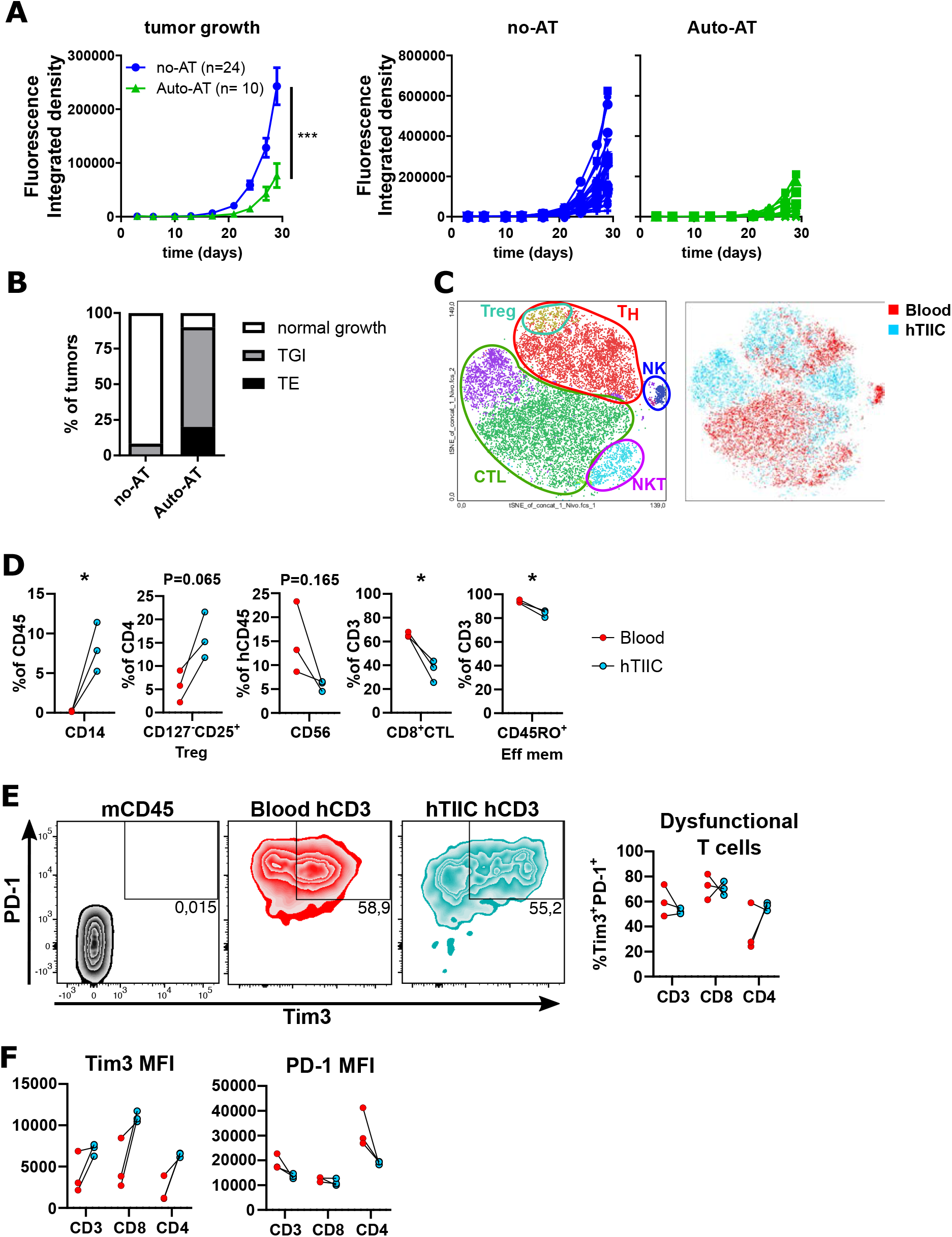
Engineered human skin fibroblast-derived tumors are recognized by autologous immune cells in Auto-AT mice. (A) Growth curves for 4T transformed adult dermal skin fibroblasts (left) and individual growth for all tumors without immune humanization (middle panel, no-AT, blue) and with autologous Hu-AT (right panel, Auto-AT, green) expressed in radiance integrated density. Shown is the mean ±SEM (B) Tumor rejection assessment at sacrifice. TE = Tumor elimination, TGI = tumor growth inhibition. (C) Characterization of the human immune infiltrate by flow cytometry. tSNE Dimensional reduction visualization with unsupervised clustering using FlowSOM module for FlowJo and manual labelling of subtypes (left). Differential clustering between hTIIC and blood human CD45^+^ cells shows little overlap signifying differential marker expression levels (right). (D) Manual quantification of differentially represented human immune populations between blood and tumor samples. (E) Exhaustion/dysfunction gating strategy (left) and quantification (right) showing no significant change in total CD3, CD8+ and CD4+ T cell population dysfunction frequency between blood and tumor. (F) Differential expression levels of dysfunction markers Tim3 and PD-1 on human T cell populations in blood vs. tumor samples shown by mean fluorescence intensity quantification. In panels (C, right), (D), (E) and (F), red = Blood human immune cells, and light blue = hTIIC. n= number of tumors, 2 tumors per mouse.

We next wanted to measure the immunogenicity of fibroblastic tumors in Hu-BLT mice which allow for a robust reconstitution of diverse functional immunological compartments. Using the same methodology used to generate 4T fibroblastic tumors derived from adult skin fibroblasts, we generated new 4T fibroblastic tumors derived from fetal skin. When injected in Hu-BLT mice, these tumors were partially or completely rejected in both allogeneic and autologous conditions (Fig. 2A). As expected, the growth of allogeneic tumors was more efficiently inhibited than the growth of autologous tumors in Hu-BLT mice (Fig. 2A and 2B). Flow cytometry analysis of the human immune compartment (Fig. 2C-D and S2) revealed no phenotypic variations in circulating human blood cells between mice harboring Auto- and Allo-tumors (Fig. 2C left and 2E). hTIIC of non-eliminated tumors displayed similar population clustering with a slight but significant enrichment of CD8^+^ cells and concomitant reduction in CD4^+^ cells in Auto-BLT tumors (Fig. 2C right and 2D-E). As observed in Auto-AT mice, major phenotypic variations between circulating and tumor-infiltrating human immune cells were observed for both Auto- and Allo-BLT mice (Fig. 2C-E). However, while hTIIC displayed a marked increase in CD4^+^CD25^+^CD127^−^Treg cells and in PD-1 and Tim3 expression, circulating T cells did not show any sign of exhaustion in opposition to what we observed in Auto-AT mice (Fig. 2D-E). Yet, most tumors were eventually able to evade the immune response in Hu-BLT mice suggesting that they can also induce an immunosuppressive microenvironment that cannot be overcome despite the apparent absence of exhaustion in circulating blood T cells. Overall, these results demonstrate it is possible to achieve partial or total immune rejection of engineered human fibroblastic tumor cells in humanized mice.

**Figure 2.**
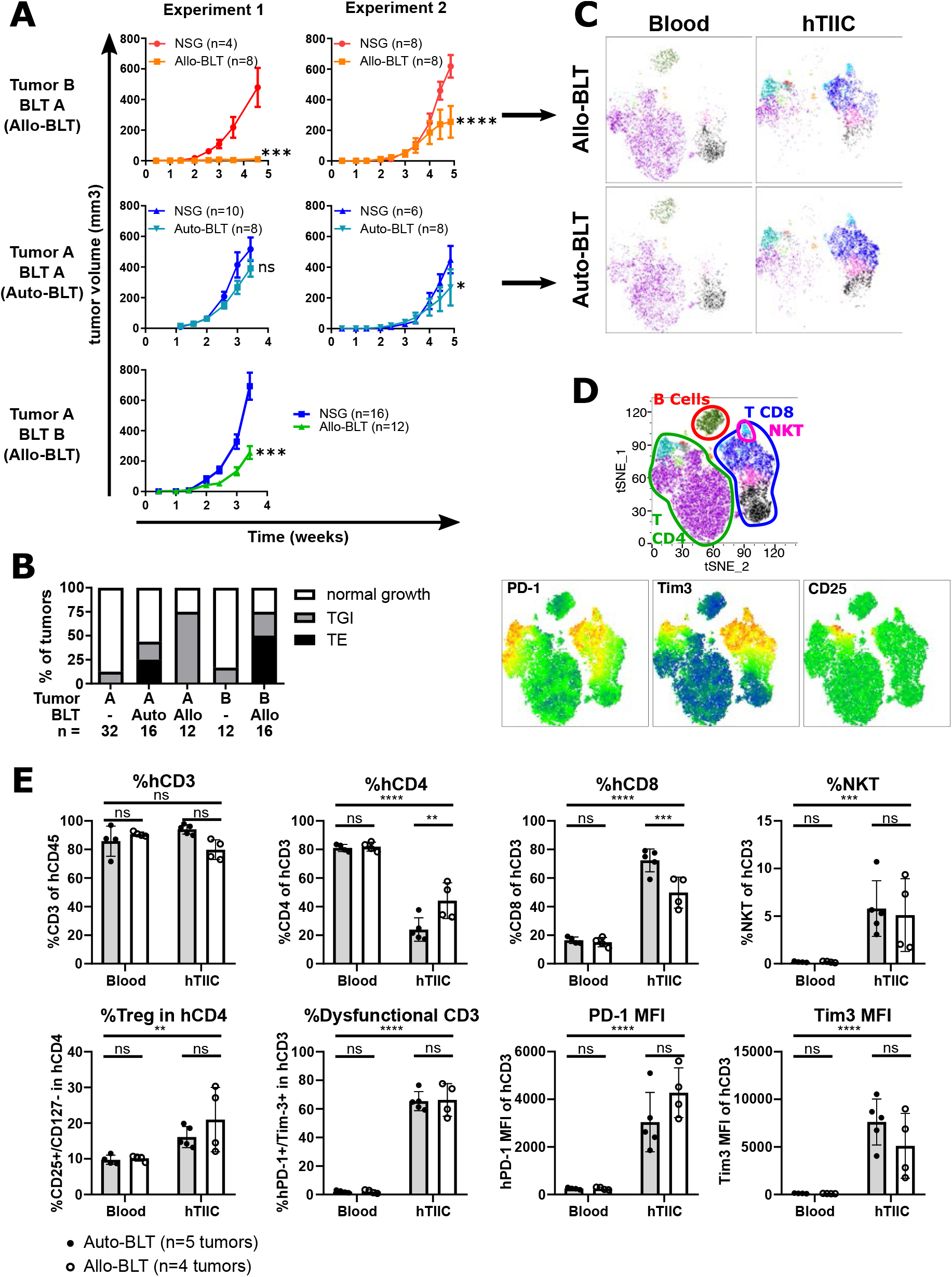
Human skin fibroblast-derived tumors are recognized by autologous immune cells in Hu-BLT mice. (A) Growth curves for repeated experiments showing fetal skin fibroblast-derived tumors from two different donors exposed to allogeneic (top and bottom) and autologous (middle) Hu-BLT immune reconstitution. Shown as mean±SEM, n= number of tumors, 2 tumors per mouse. (B) Tumor rejection assessment at sacrifice for each condition presented in (A). TE = Tumor elimination, TGI = tumor growth inhibition showing Auto-BLT to be less proficient at rejecting tumors than Allo-BLT. (C) tSNE dimensional-reduction plots of human blood (Left) or tumor infiltrating immune cells (hTIIC, right) for Allo-BLT (top) and Auto-BLT (bottom) flow cytometry samples. All immune cells are from the same donor. (D) Population annotation of human immune populations in BLT-humanized mice. Combined results and population annotation from panel (C) (top) and expression of exhaustion markers PD-1 and Tim3 (bottom left and center) and Treg-associated marker CD25 (bottom right). All these markers are enriched in hTIIC samples. (E) quantification of effector populations (top row) in blood and hTIIC samples of Auto-BLT (grey bars, full circles) and Allo-BLT (empty bars and circles) samples. Enrichment of CD8^+^ cells and concomitant decrease in hCD4^+^ T cells in Auto-BLT was observed. Quantification of immunosuppressive Treg (bottom far left) and dysfunctional T cells (bottom center-left), and expression levels of PD-1 (bottom center-right) and Tim3 (bottom far right). No significant variation between immunosuppressive and dysfunction markers between Auto- and Allo-BLT samples was observed.

### Tumors originating from transformed iPSC-derived hepatic cells are recognized by autologous immune cells in Hu-AT mice

To generate alternative tumor models that can be used in a humanized setting, we set out to transform iPSC-derived cells from healthy donors. First, iPSC were differentiated into hepatocytes using a previously described protocol (schematized in Fig. 3A) (24). We initiated the transformation process using SV40ER at different timepoints in the cell differentiation process which led to the formation of growing colonies. These colonies were then transduced with HRas^V12^ and hTERT lentiviral particles after the differentiation protocol was complete and then finally modified to express the firefly luciferase marker (Fig. 3A). We tried transforming cells at day 16, 22 and 30 of the differentiation protocol and observed the transformation potential of the cells was overall very limited. Indeed, only few colonies (< 10) were formed in day 16-transduced cells and only rare colonies emerge when cells were transduced on day 22. It was not possible to transform cells at day 30, when hepatocytes reach a quiescent state, suggesting that *in vitro* transformation using SV40ER is possible only in a subset of yet not fully differentiated cells and likely requires cells to maintain a proliferation potential.

**Figure 3.**
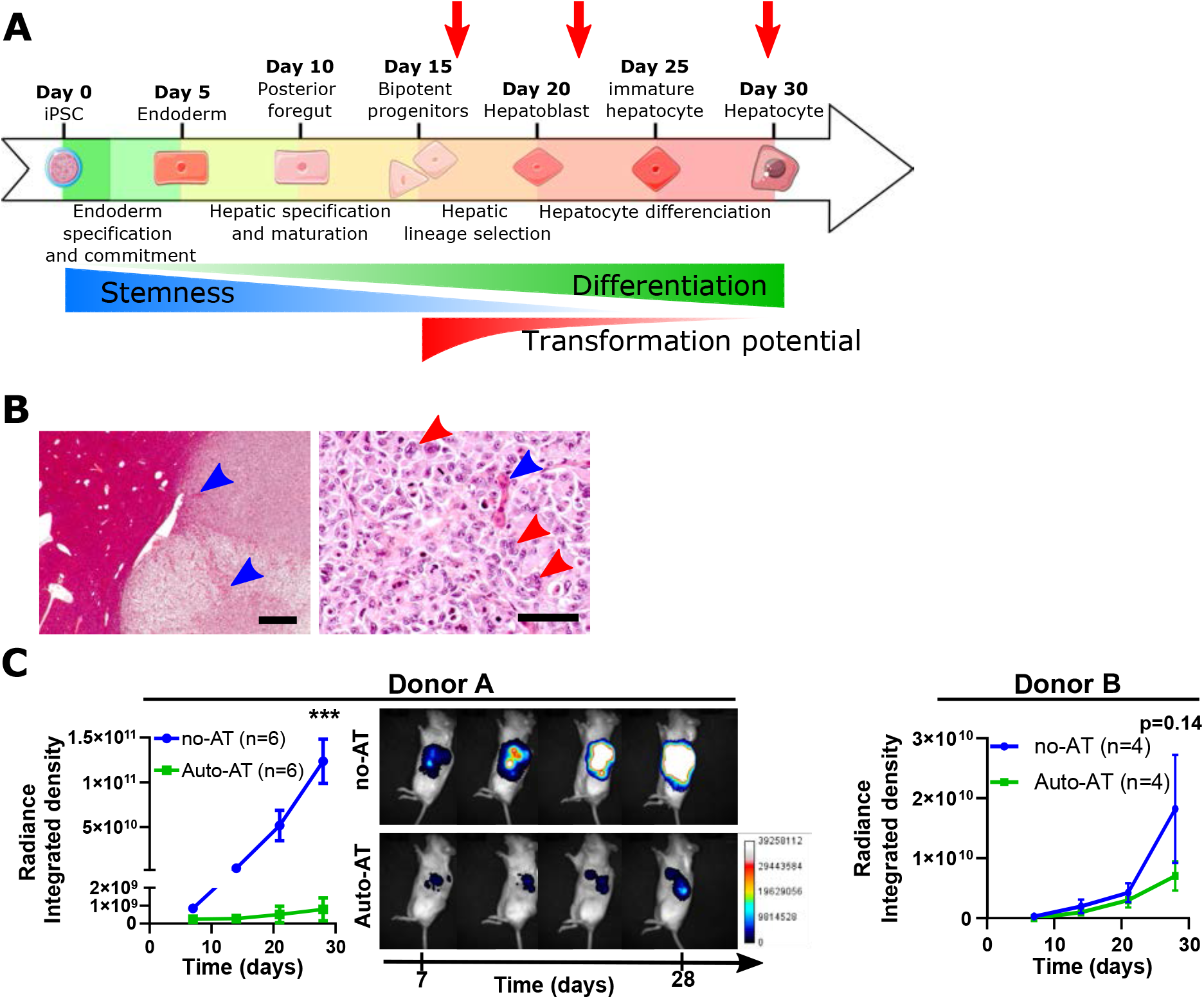
iPSC-derived hepatic tumors are recognized in Auto-AT mice. (A) Schematic of the iPSC differentiation protocol used to generate hepatocyte-like cells (HLC). Red arrows indicate timepoints for initiation of cellular transformation by SV40ER transduction. (B) Histology of an HLC 4T tumor at low and high magnifications of hematoxylin eosin saffron (HES) staining. High magnification (left) shows border of well-circumscribed tumor with entrapped liver parenchyma (blue arrowheads) and varying tumor density. High magnification photomicrograph (right) again shows parenchymal entrapment (blue arrowheads), polynucleated cells (red arrowheads), numerous mitoses, hyperchromatic nuclei, and generally highly pleomorphic cells and nuclei. (C) Mean±SEM of *in vivo* HCT 4T tumor elimination by Auto-AT. Integrated density of intrahepatic tumor-associated luciferase signal for two independent donors (left and right) and representative longitudinal *in vivo* bioluminescence from donor A. Scale bar in C, left = 1 mm and right = 100 μm.

Characterization by qPCR of untransformed hepatocyte-like cells (HLC) showed these cells express liver-specific markers (alpha fetoprotein (AFP), hepatocyte nuclear factor 4 alpha (HNF4A), albumin (Alb) and asialoglycoprotein receptor 1 (ASGR1)) at generally comparable levels compared to the control HepG2 hepatocellular carcinoma cell line (Fig. S3). Upon transformation of HLC (named HLC 4T), the expression of most markers was decreased (except for the bipotent hepatoblasts and cholangiocyte-associated cytokeratins 19 and 7) suggesting either that transformation induced dedifferentiation of the cells or that transformation occurred more efficiently in less differentiated progenitors. When injected intrahepatically (1 and 5×10^5^ cells for donor A and donor B respectively), HLC 4T cells formed circumscribed tumors with entrapped liver parenchyma and mixed sarcomatoid components (Fig. 3B left). Tumor cells were highly mitotic, undifferentiated, pleomorphic and occasionally hyperchromatic with some polynucleated cells and ill-defined cell borders (Fig. 3B right). AFP staining was weakly positive *in vitro* when compared to the control HuH-6 hepatoblastoma cell line but negative *in vivo* (Fig S3, top), Alb expression was strongly positive *in vitro* and weakly positive *in vivo* (Fig. S3, center) and HepPar1 staining was negative *in vitro* and *in vivo* (Fig. S3, bottom). Pathological examination suggests that HLC 4T tumors resemble undifferentiated embryonic sarcoma of the liver (25). Interestingly, as observed with fibroblastic 4T cell lines, the intrahepatic injection of HLC 4T cells formed tumors in all animals with similar kinetics. The addition of 5×10^6^ autologous PBMCs and granulocytes was enough to partially or completely reject HLC 4T tumors in NSG-SGM3 mice in two independent donors (Fig. 3C). These results suggest it is possible to generate hepatic-like tumors and that these tumors are immunologically detected in Auto-AT mice.

### iPSC-derived neural tumors recapitulate high grade glioblastoma and are recognized by autologous immune cells in Hu-AT mice

To further explore the flexibility of our models we initiated the transformation of iPSC-derived neural stem cells (iNSC) and astrocytes (iAstro). Cells were first differentiated using an establish protocol (see Fig. 4A and methods) and then characterized for their expression of key differentiation markers (Fig. S4). The transformation of iNSC and iAstro (named iNSC 4T and iAstro 4T) was achieved using the same set of oncogenes described previously and initiated at a single time point immediately upon confirmation of the cells acquiring the nestin and glial fibrillary acidic protein (GFAP) markers respectively. Transformed cells led to the generation of highly aggressive tumors when injected orthotopically in NSG-SGM3 mice (Fig. 4B). As few as 1.5×10^3^ iAstro 4T cells were enough to consistently generate tumors within 3-4 weeks. iNSC 4T tumors also formed highly undifferentiated tumors in NSG-SGM3 mice (Fig. 4B top), however the histopathology was inconclusive in characterizing them as classical glioblastomas multiform (GBM) (Fig S5). They were nonetheless highly proliferative tumors with prominent growth along Virchow-Robin spaces but with a limited diffuse infiltration and a distinctive accumulation near the dentate gyrus in all animals (Fig 4B top). In contrast, iAstro 4T tumors were also highly proliferative but displayed a more conspicuous diffuse infiltration, parenchymal entrapment and ventricular and leptomeningeal dissemination along with extensions in the Virchow-Robin space. Variable anaplastic features and a significant involvement of reactive astrocytes were also observed (Fig 4B middle and bottom). Morphologically, iAstro 4T tumors were polymorphic and harbored multiple characteristic structures such as myxoid material, pre-necrotic cells, karyorrhexic nuclei, cellular monstruosities/giant cells and some regions akin to giant cell and epithelioid GBM. Immunophenotypically, iAstro 4T tumors were strongly vimentin- and moderately cytokeratin AE1/AE3-positive (Fig. S6 left column) and lacked expression for MAP2 and S100 (Fig. S6). Olig2, CD56, desmin and cytokeratin 19 were also found to be negative in these tumors (data not shown). Consistent with a previous study using a similar transformation approach, *in vivo* tumors lacked GFAP expression despite *in vitro* expression (26)(Fig. S5 and S6). As observed with our other tumor types, iAstro 4T tumors from two independent donors were partially or fully rejected by Auto-AT (Fig. 4D). In Auto-AT mice, both iAstro 4T and iNSC 4T orthotopic tumors were highly infiltrated by hCD45 cells, mostly CD8 T cells (Fig. 4D and Fig. S5B). Little to no infiltration was observed in surrounding mouse tissue, suggesting a specific T cell-mediated anti-tumor response. These results demonstrate that it is possible to transform iPSC-derived neural cell lines and that these cells are infiltrated by autologous immune cells upon orthotopic injection in the brain of Hu-AT mice.

**Figure 4.**
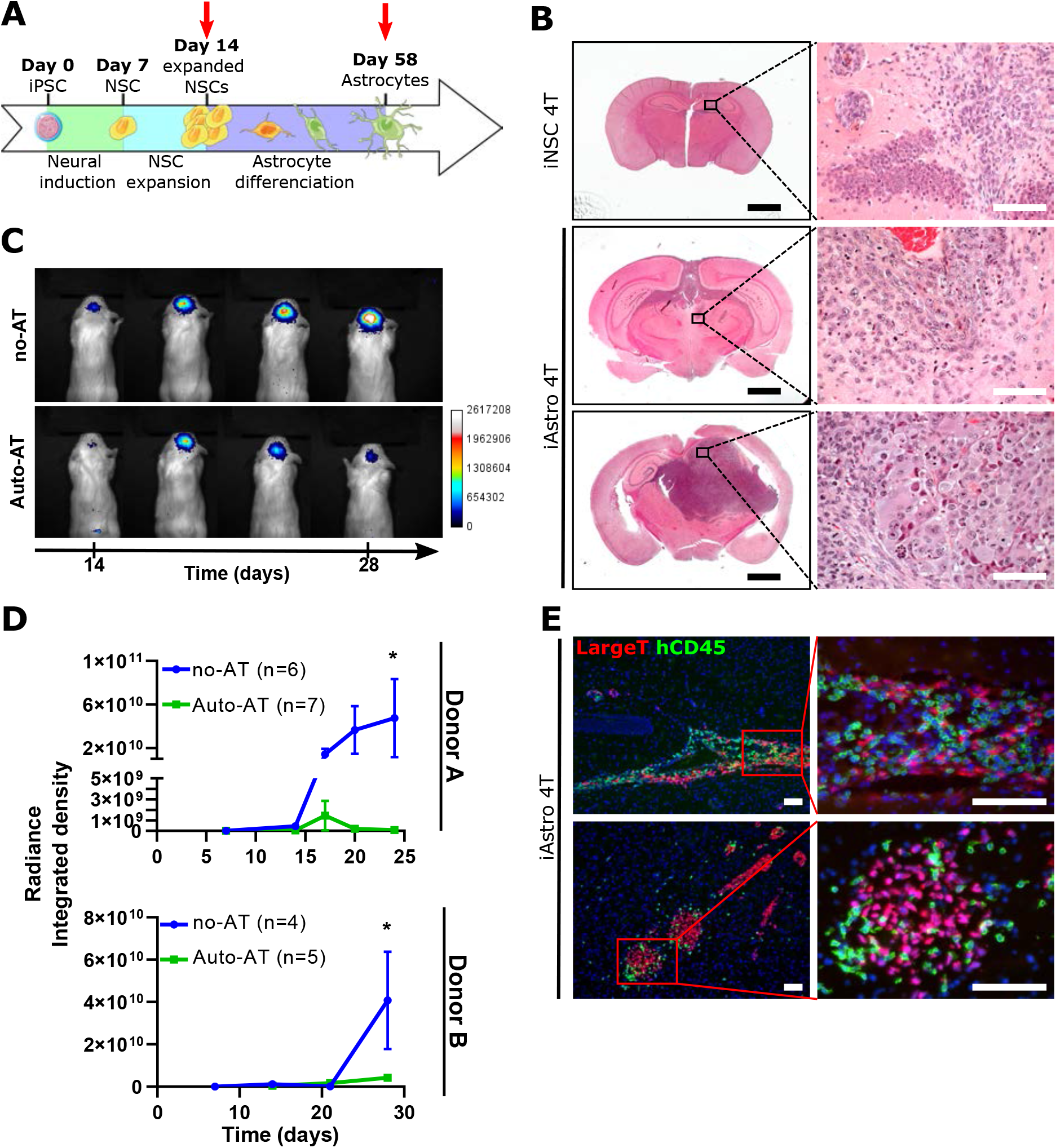
iPSC-derived neural tumors are rejected in Auto-AT mice. (A) Schematic of iPSC differentiation approach for the generation of neural stem cells (NSCs) and astrocytic cell populations. Red arrows indicate populations transformed using the 4T approach. (B) Histology of one iNSC 4T tumor (top) and two representative iAstro-derived tumors (middle and bottom). High magnification photomicrographs on right show poorly differentiated tumor cells with brisk mitotic activity with little (top) or more conspicuous (middle) diffuse infiltration, or epithelioid/giant cell differentiation (bottom). (C) Representative images of longitudinal in vivo luciferase imaging in no-AT (top) and Auto-AT (bottom) mice. (D) Mean±SEM graph of *in vivo* luciferase signal quantification of iAstro 4T tumors with (Auto-AT) and without (no-AT) adoptive transfer in two different donors. (E) Immunofluorescent staining images for human immune cells infiltrate detection within samples of iAstro 4T tumors at day 29 post tumor cell injection and Auto-AT showing human immune infiltrate specifically within tumors. Red = SV40 Large T, Green = hCD45, Blue = DAPI. Scale bar in B, left = 2 mm, all other scale bars = 100μm.

### Anti-PD-1 immunotherapy in Hu-AT mice leads to increased immune infiltration and clearance of autologous tumors

Based on the observation that our 4T tumors expressed high levels of PD-L1 (Fig. 5A), we next investigated if Hu-AT mice with autologous 4T tumors would represent a good model to evaluate the efficacy of anti-PD-1 immunotherapy. Our results showed that treatments of Hu-AT mice with the blocking antibody Nivolumab lead to a slight, but not significant, reduction in the size of fibroblastic tumors (Fig. 5B). Yet, flow cytometry analysis showed a significant increase in the relative abundance of human CD45^+^ cells infiltrating tumors in Nivolumab-treated animals (Fig. S7), an observation we also confirmed by immunofluorescent staining (Fig. 5D). However, further analysis revealed that Nivolumab did not significantly affect immune population profiles either in blood or within tumors as shown by highly overlapping populations in tSNE dimensionality-reduction plots (Fig. S7). Only CD3^−^CD56^+^ NK cells were slightly decreased in abundance in blood samples of Nivolumab-treated animals (Fig. S7A), and nearly all human infiltrating immune cells were CD3^+^ T cells (Fig S7B). No specific population enrichment was observed within the hTIIC compartment in Nivolumab-treated animals. Phenotypically, infiltrating T cells were also largely similar between animal groups with only PD-1 staining varying significantly, presumably as a staining artifact due to steric interference by residual circulating Nivolumab. Because fibroblastic tumors proved to be generally more difficult to reject in comparison to HLC and iAstro tumors (compare Fig 1A with Fig. 3C or 4D), we repeated the injection of Nivolumab in mice with HLC 4T tumors. In conditions where these tumors were not fully rejected in Auto-AT mice, treatment with Nivolumab was able to significantly improved tumor rejection (Fig. 5C). Because HLC 4T tumors from Nivolumab injected mice never reached a size that would have permitted their surgical retrieval, it was unfortunately not possible to analyze the tumor immune infiltrate in this model. Overall these results suggest the injection of Nivolumab in mice harboring PD-L1+ engineered autologous tumors can increase the infiltration of T cells and delay the growth of certain tumor types.

**Figure 5.**
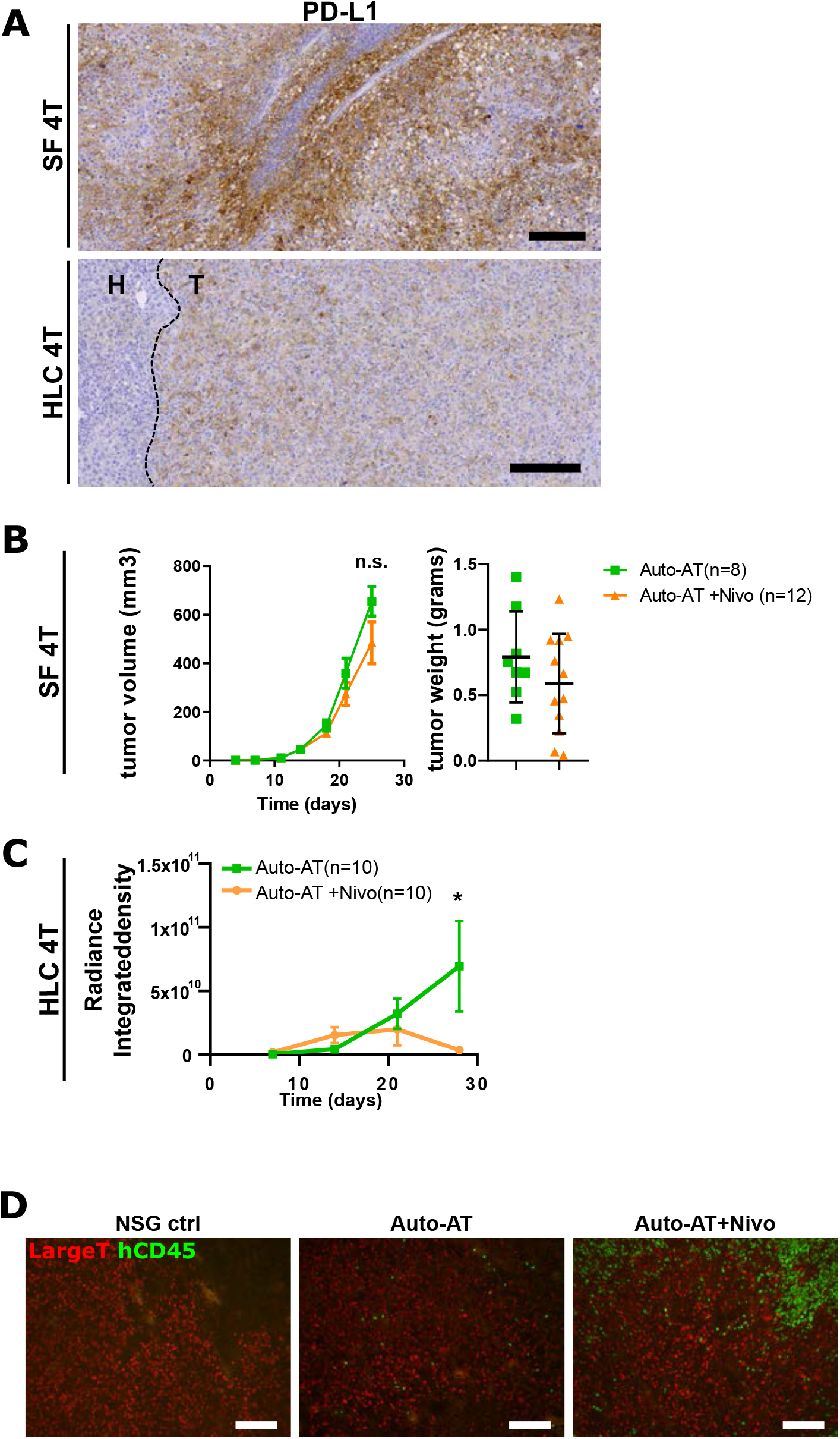
Injection of Nivolumab in Hu-AT mice leads to increased immune infiltration and clearance of autologous tumors. (A) Tumor microphotograph of PD-L1-stained fibroblastic (top) and HLC 4T (bottom) tumors. Fibroblastic tumors show strong PD-L1 staining and HLC 4T tumors show tumor-specific (T) staining when compared to surrounding mouse hepatic tissue (H). (B-C) Effect of Nivolumab administration in Auto-AT fibroblastic (B) and HLC 4T (C) tumor-bearing mice. (B) Shown are the results of 2 independent experiments showing tumor growth (left panel, Mean±SEM) and tumor weight at sacrifice (right, Mean±SD) for each experiment. n= number of tumors, 2 tumors per mouse. (C) Tumor growth curve (Mean±SEM) of HLC 4T tumors as measured by luciferase-associated radiance in no-AT (blue), Auto-AT (green) and Auto-AT (orange) treated with Nivolumab at days X, Y and Z (red arrows). n = number of mice, 1 intrahepatic tumor injection per mouse. (D) Representative images of human immune infiltrate within s.c. fibroblastic (top) and HLC 4T (bottom) tumor samples for NSG-SGM3 control (left), Auto-AT alone (middle) and Auto-AT +Nivolumab (right). Red = Large T, green = human CD45, Scale bars = 200μm.

## DISCUSSION

In this study, we generated genetically-defined tumor cell lines from primary and iPSC-derived cells that, when combined with immune humanization of mice, can be used for the evaluation of cancer-immune cell interactions in an autologous setting. In comparison to patient-derived cancer cell lines or xenograft, our 4T tumors are more easily accessible, genetically homogeneous, can be transformed using a customizable process and have not undergone immunoediting events that can interfere with primary immune rejection. While this represents a limitation for studying acquired immune resistance mechanisms, it can be advantageous for whoever is interested in screening for such mechanisms or validating new targets in a genetically defined environment. The versatility of the model also allowed us to compare the efficacy of Nivolumab against different tumor cell types. Indeed, the strong expression of PD-L1 in our 4T tumors was not surprising given RAS signaling was shown to stabilize PD-L1 mRNA (27). Yet, we also unexpectedly observed that iNSC and iNSC 4T tumors acquired the expression of the disialoganglioside GD2, a tumor-associated antigen (Fig.S8). This example suggests that this tumor model could also be used for the testing of GD2-autologous chimeric antigen receptor T cells therapies.

Overall, we observed that fibroblastic tumors were less efficiently rejected compared to iPSC-derived hepatic and neuronal tumors. Beside the fact that these are intrinsically different tumors, other factors may explain this difference. We believe the subcutaneous injection of fibroblastic tumors, as opposed to intra-hepatically or intracranial, likely limited the early access to immune cells allowing the tumor to grow unchallenged, at least in the first few days/weeks. In support of this, we noticed HLC-4T tumors were less efficiently rejected when injected subcutaneously compared to inta-hepatically (data not shown).

Here, we choose to work with an aggressive transformation approach (SV40ER/HRas^V12^/hTERT) which systematically lead to fast-growing, aggressive and highly undifferentiated tumors. Yet, we noted the inability of this approach to transform quiescent cells *in vitro*, such as terminally differentiated HLC and GFAP-rich iAstro cells. Indeed, we observed that only experimental conditions containing residual proliferating cells were permissive to transformation by our oncogenes. This is in opposition to still proliferating fibroblasts and NSC cultures which were more uniformly transformed. While this could be an artefact of non-physiologic cell culture conditions, this observation supports the hypothesis that most cancer cells originate from stem or progenitor cells rather than terminally differentiated quiescent cells (28). In support with the cancer stem cell hypothesis, we noticed that transformed iAstro 4T cells expressed the glioblastoma cancer stem cell associated marker CD133 in vitro (29) (Fig S9). Going forward, we propose that alternate tailoring of driver genetic alterations could improve different aspects of tumor phenotype and maintain tumor cells at a higher level of differentiation. For examples, driver mutations such as IDH1, ATRX and EGFR in glioblastoma or CTNNB1, NFE2L2, APOB and ALB in hepatocellular carcinomas could help steer these models towards the desired histopathology (30). In the future, more adapted transformation protocols combined with the development of cancer organoid may be more representative of spontaneously emerging human tumors (31). Moreover, the possibility to combine gene editing tools to recapitulate precise somatic mutations with novel humanized mouse models resistant to GvHD, should also allow for the development of more faithful iPSC-derived cancer models (31–33). This is particularly important given T cells in our AT-mice showed activation/exhaustion profiles suggesting that the mice were at an early stage of GvHD which prevented us from tracking tumor growth long-term. Using newly developed MHC knockout mouse strains should help circumvent this problem in the future (33,34).

Of note, is the fact that all of our cell lines generated with the 4T transduction approach were immunogenic to various degrees *in vivo*. One possibility is that non-human proteins such as the viral SV40 large and small T antigens, which express immune-dominant epitopes are mainly recognized (35). However, it is also possible that such a broad transformation approach, which systematically lead to poor differentiation phenotypes, as demonstrated with our HLC and iAstro 4T tumors, induces immunogenic tumor antigens, or that partially differentiated cells maintained the expression of embryonic antigens previously shown to be able to activate an immune response (36). Our models were effective in mounting an autologous immune response mediated mostly by effector T cells, as other effector types are poorly reconstituted in most humanized models. NK cells, for instance, require exogenous (37) or transgenic addition of IL-15 (38) or IL-2 (39) for their development and maturation in HSC-reconstituted mice. In this study, however, we suspect NK cells to be non-essential to tumor recognition and elimination for few reasons. First, while CD3^−^CD56^+^ NK cells were identified in some Auto-AT animals, not surprisingly none were detected within tumors in absence of cytokines supporting NK cell expansion. Also, the inability of BLT mice in the NSG background to reconstitute functional NK cells (40) did not prevent them from mounting an anti-tumor response similar to what we observed in Auto-AT mice. Finally, NK cells purified from peripheral blood failed to show a cytotoxic response against fibroblastic tumors *in vitro* (data not shown).

In conclusion, we generated highly accessible and flexible autologous models of tumor-immune cell interactions which should facilitate the evaluation and development of cancer immunotherapies. We foresee that ongoing development in our ability to adequately differentiate human iPSC into a variety of cell types and organoid, including hematopoietic stem/progenitor cells for immune reconstitution in mice, will open many more possibilities in a near future.

## ACKNOWLEDGMENTS

We are grateful to the flow cytometry and animal facility for providing technical support and to Renée Dicaire for handling clinical samples.

## AUTHOR CONTRIBUTIONS

G.M.B., B.B., D.M., O.L., Y.L., M-A.M., and C.C., performed experiments. C.R., M.P., E.H. provided material, reagents and expertise. D.D.S, B.E, provided expertise, J.V.G. provided material, G.M.B. and C.B. designed the studies, G.M.B. and C.B. wrote the manuscript.

## METHODS

### Isolation and culture of skin fibroblasts

Skin fibroblasts were isolated from either adult skin biopsies or fetal skin segments for Hu-AT or Hu-BLT models respectively. Skin tissues were obtained from consented healthy adult donors or after surgical abortion at around week 20 of pregnancy in accordance with the Bureau d’Éthique à la Recherche du CHU Sainte-Justine (protocol number 2017-1476). In both cases, the tissue was cleaned out to preserve only the dermis and epidermis, triturated into 1-5mm^2^ pieces and digested with collagenase D (Roche) for one hour at 37°C with agitation. The whole mixture was then centrifugated at 400 g for 5 minutes and washed with DMEM (Wisent) twice. The whole product of digestion was seeded in T150 flasks in DMEM with 10% FBS and 0.2% primocin (Invivogen). Subsequent passages were also maintained in DMEM with 10% FBS and 0.2% primocin.

### iPSC reprogramming

Heathy human donor PBMCs or skin fibroblasts were reprogrammed into iPSCs using the integration-free based Sendai virus (Cytotune 2.0 kit from Life Technologies). Low passage primary fibroblasts were used to increase reprogramming efficiency. Following transduction, emerging clones were manually picked and cultured under feeder-free conditions using Geltrex-coated dishes and Essential 8 medium (Life Technologies). iPSC clones were maintained in culture for at least 15 passages to ensure stable pluripotency before characterization was conducted by the iPSC – cell reprogramming core facility of CHU Sainte-Justine. Cells were shown to have a normal karyotype and colony to express the human SSEA-4, Sox2, OCT4 and TRA1-60 makers (pluripotent Stem Cell 4-marker Immunocytochemistry Kit from Life Technologies as shown previously)(41).

### Hepatocyte differentiation

Differentiation of human iPSCs into hepatocyte-like cells (HLC) was achieved according to our 30-day long, previously described protocol mimicking liver development (24). In brief, iPSCs were differentiated on laminin-coated plates through consecutive stages (primitive streak, mesendoderm, definitive endoderm, posterior foregut, hepatoblasts and, eventually, HLC). Differentiation was started by changing the medium to RPMI-B27 minus insulin (Life Technologies) supplemented with 1% KOSR (Life Technologies) and 100 ng/ml Activin A (R&D Systems) for 3 days. For the first 2 days cells were also exposed to 3 μM CHIR-99021 (Stem Cell Technologies). From day 4 for 5 days, RPMI-B27 (minus insulin) medium was supplemented with 20 ng/ml BMP4 (Peprotech), 5ng/ml bFGF (Peprotech), 4 μM IWP-2 (Tocris), and 1 μM A83-01 (Tocris). From day 10 to day 15, the medium was changed to RPMI-B27 supplemented with 2% KOSR, 20 ng/ml BMP4, 5 ng/ml bFGF, 20 ng/ml HGF (Peprotech) and 3 μM CHIR-99021. At day 16, the medium was changed to HBM/HCM (minus EGF) medium (Lonza), supplemented with 1% KOSR, 20 ng/ml HGF, 20 ng/ml BMP4, 5 ng/ml bFGF, 3 μM CHIR-99021, 10 μM dexamethasone (Sigma), and 20 ng/ml OSM (R&D System) for 5 days. From day 20, for 5 days, HBM/HCM medium was supplemented with 1% KOSR, 10 μM dexamethasone and 20 ng/ml OSM. From day 25, the cells were maintained in HBM/HCM 1% KOSR medium supplemented with 10 μM dexamethasone. Medium was changed every day for the first 20 days and every other day from day 20 to 30. During all the differentiation process, the cells were kept at 37°C, ambient O_2_ and 5% CO_2_.

### NSC and Astrocyte differentiation

Neural Stem Cells (NSC) and Astrocytes were differentiated from hiPSCs by first using the PSC Neural Induction Medium (NIM) (Gibco). Briefly, hiPSCs were dissociated into small colonies using the gentle cell dissociation reagent (Gibco) and seeded on Geltrex matrix (Gibco) at 50% confluence in E8 medium. 24h later, E8 medium was replaced with complete NIM (neurobasal medium with neural induction supplement) for 7 days with medium change at days 2, 4 and 6. Resulting Neural stem cells (NSCs) were then expanded for at least 3 passages in neural expansion medium (neurobasal medium/ advanced-DMEM/F12 with neural induction supplement) (Life Technologies). Resulting cells were considered NSCs after validation of the acquisition of nestin expression and loss of OCT4 pluripotency marker by immunofluorescence. NSCs were maintained in StemPro NSC SFM for further transformation (see below) or differentiated further into the astrocyte lineage. To do so, NPCs were plated on Geltrex in a 6 well plate in StemPro NSC SFM supplemented with 20ng/ml of EGF and bFGF for 2 days before switching to astrocyte differentiation medium consisting of DMEM with 1% N-2 supplement (Gibco), 1% GlutaMAX-I supplement (Gibco) and 1% FBS (Wisent) with medium change every 3-4 days. Differentiated astrocytes were typically observed on days 5-7. Astrocytes were characterized *in vitro* by immunofluorescence for the expression of GFAP and Vimentin. Briefly, cells were seeded in Geltrex coated Lab Tek chamber slides (Nunc) and fixed 5 days later in 3.7% formaldehyde before permeabilization and blocking using PBS with 0.3M Glycine (Wisent), 1% BSA (Sigma), 2.5% Goat Serum (Sigma) and 0.1% Triton X-100 (Sigma) for 30 minutes at room temperature. Primary anti-GFAP (Dako) or anti-Vimentin (BioLegend) antibodies were used at 1/200 concentration overnight at 4°C and secondary AF488-conjugated anti-mouse Ig (Life Technologies) at 1/500 for 1 hour at room temperature. GFAP-expressing astrocyte populations were subsequently transformed (see below). Flow cytometry was done using anti-CD133/1-PE (Miltenyi Biotec, clone AC133) and anti-GD2-APC (Biolegend, clone 14G2a).

### Lentivirus production

All lentiviral particles were produced by overnight PEI transfection of 293T/17 cells (ATCC CRL-11268) in complete RPMI with 10% FBS with 2^nd^ (pPAX2) or 3^rd^ (pMDL and pRSV-Rev) generation packaging plasmids along with a plasmid encoding the VSV-G envelope. SV40ER was subcloned from pBABE SV40ER from William Hahn (Addgene #10891) into a lentiviral transfer plasmid containing a Neomycin resistance gene (SV40ER-Neo), RasV12 lentiviral transfer plasmid containing a puromycin resistance gene (RasV12-puro) was obtained from Francis Rodier (CHUM, Université de Montréal), hTERT lentiviral transfer plasmid was generated as previously described (42), mPlum was subcloned from pQC mPlum XI from Connie Cepko (Addgene #37355)(43) into a lentiviral transfer plasmid containing puromycin selection gene and firefly luciferase IRES-GFP (luc/GFP).

### Cellular transformation

Primary fibroblasts and astrocytes (ScienCell research Laboratories, CA USA) or iPSC-derived cells lines were transformed with lentiviral particles in a sequential order. For fibroblast-derived tumors, cells were transduced overnight with medium containing SV40ER-Neo viral particles. Three days later, G418 selection was added at 300 μg/mL (ThermoFisher) and maintained until control GFP-transduced cells were eliminated. Cells were subsequently transduced with RasV12-puro lentivirus in a similar way. Puromycin (ThermoFisher) selection was started three days post-transduction at 2 μg/mL again with GFP-transduced cells as control. Cells were then transduced overnight with medium containing hTERT lentiviral particles. 2 days later, cells were transduced overnight with mPlum viral particles. All transductions were done in presence of 8 μg/mL polybrene. Cells were subsequently expanded and sorted on FACSAriaII (BD Biosciences) in the APC channel for the expression of mPlum. For hepatocyte, astrocyte and NPC-derived tumor cells, lentiviral suspensions were used as well as luciferase/GFP instead of mPlum. However, the initial overnight SV40ER transduction was made at day 16, 22 or 30 of the hepatocyte differentiation protocol (see above), and differentiation was pursued. Cells were passaged on day 32 and RasV12-puro transduction was done once the cells were 70-80% confluent. Three days later, concomitant G418 and puromycin selection started, still at 300 μg/mL and 2 μg/mL respectively, until individual controls for each antibiotic were eliminated. Cells were subsequently transduced with hTERT then luciferase/GFP and sorted on FACSAriaII (BD Biosciences) for GFP expression.

### Mouse reconstitution

All *in vivo* experiments were conducted in conformity with institutional committee for good laboratory practices for animal research (protocol #669). Male and female mice used in this study (aged between 7-18 weeks) were of the NSG and NSG-SGM3 (expressing human IL3, GM-CSF and SCF) background. All mice were breed onsite and housed in the animal facility at the CHU Sainte-Justine Research Center under pathogen-free conditions in sterile ventilated racks after being originally obtained from the Jackson Laboratory (Bar Harbor, ME). For adoptive transfer (Hu-AT) experiments, human adult peripheral blood was harvested from healthy donors after informed consent and immune cell isolation using Ficoll-Paque gradient (GE Healthcare). Buffy coat was harvested for PBMCs while granulocytes were isolated from the gradient pellet: Ficoll-Paque was aspirated and the pellet was broken and resuspended in 38 mL of sterile deionized water for 20 seconds for RBC lysis before addition 2 mL of sterile 20X PBS solution. PBMCs and granulocytes were counted and mixed at 1:1 ratio before injection into mice. 5×10^6^ PBMCs and 5×10^6^ granulocytes for a total of 1×10^7^ WBC were injected intraperitoneally (i.p.) into NSG-SGM3 in 200 μL total volume. Aged-match NSG-SGM3 without i.p. injections of WBC were used as no-AT controls.

For BLT-reconstituted mice (Hu-BLT), 6-8-week-old NSG mice were sublethally irradiated with 2 Gy total body irradiation using a Faxitron CP-160 before surgical implantation of 1-2mm^3^ human fetal thymus under the kidney capsule and intravenous injection of 5×10^5^ CD34+ hematopoietic stem cells (HSC) isolated from autologous fetal liver as previously described (44). Fetal (16-21 weeks) tissues were obtained after written informed consent (ethical committee of CHU Sainte-Justine, CER#2126). Hematopoietic engraftment was assessed by flow cytometry at 4 and 8 weeks by staining 50 μL of peripheral blood collected from the saphenous vein. The following antibodies were used (mouseCD45-FITC from BD bioscience and 7-AAD, humanCD45-PE/Cy7, humanCD19-PE, humanCD3-APC and humanCD14-APC/Cy7 from Biolegend) and cells analyzed on the LSRFortessa flow cytometer (BD Biosciences). Only mice with high reconstitution at week 8-10 (35-75% human CD45, >20% CD3) were used in this study. Age and sex-matched non-reconstituted NSG mice are used as negative control. In all cases, mice showing signs of GvHD were removed from analysis.

### Mouse orthotopic injections and monitoring

All injections and surgical procedures were undergone under aseptic conditions in the CHU Sainte-Justine animal facility. For the subcutaneous injections of fibroblast-derived tumors, 5×10^5^ cells were injected in 100 μL of RPMI medium on each flank of isoflurane-anesthetized and previously shaved mice. *In vivo* growth monitoring was done twice weekly on the Q-Lumi *In Vivo* imaging system (MediLumine, Montreal) by fluorescent tracking of mPlum-expressing tumor cells. Fluorescence signal was standardized internally for each picture and normalized for defined parameters using FIJI macros for picture processing. Tumor signal analysis was measured semi-manually also using FIJI macros and expressed in fluorescence integrated density or by caliper measurements. For checkpoint inhibitor studies, three 6 mg/kg doses of Nivolumab anti-human PD-1 antibody (graciously offered by Bristol-Myers Squibb, New York) were injected i.p. at day 14, 17 and 21 post tumor cell injection. Tumors were excised upon sacrifice and tumor weight and volume were recorded and compared to control tumors in each experiment. Normal growth was defined by tumors bigger than one standard deviation (SD) below control mean. Tumor growth inhibition (TGI) characterized palpable, harvestable tumors smaller than one SD below control mean. Tumors were considered eliminated (TE) when unpalpable or too small to be harvested at sacrifice Mice were sacrificed when tumors reached a maximum of 1500 mm^3^ or showed signs of distress.

Intracranial injections of astrocyte- or NPC-derived tumors were done using a stereotaxic apparatus (Stoelting). Briefly, mice were anesthetized with 2.5% isoflurane and the cranium exposed by performing a 5-6 mm incision on the scalp. A burr hole was made 2mm posterior and 1mm right of the bregma. A 10 μL Hamilton syringe was then inserted into the hole at a 3mm depth and 1 μL of tumor cell suspension (containing 1.5×10^3^ cells) was slowly injected (over a period of 10 seconds). The syringe was maintained for one minute in the hole before being slowly withdrawn to avoid gushing. The hole was then closed with Vetbond tissue adhesive (3M) before suturing the scalp. Mice were treated with buprenorphine daily for 2 days following surgery and monitored for distress signs. *In vivo* monitoring of tumor growth was done at regular intervals by bioluminescence tracking of firefly luciferase-expressing tumor cells. To do so, a 30 mg/mL solution of D-luciferin (PerkinElmer 122799) was injected i.p. at a dose of 150 mg/kg and imaged after 10 minutes without filters on the Q-Lumi *In Vivo* Imaging System. Signal normalization and analysis was done automatically for all time points using FIJI macros and expressed in radiance (photons · s^−1^ · sr^−1^ · cm^−2^) integrated density (Area · mean intensity).

Intrahepatic injections of iPSC-derived hepatic cells were done as follows: Briefly, mice were anesthetized with 2.5% isoflurane and a 15 mm incision was made underneath the ribcage on the left ventral flank. A small incision was made in the peritoneal lining to expose the left hepatic lobe. Using a glass capillary, a 10 μL (containing 1 or 5×10^5^ cells for donor A and donor B respectively) injection was made at about 2 mm depth in the liver lobe. Incisions were sutured, and the animal was treated with buprenorphine daily for 2 days following procedure. *In vivo* monitoring of tumor growth was done at regular intervals by bioluminescence tracking of firefly-luciferase expressing tumor cells as described above.

### Histological analysis and staining

Whole organs or tumor tissues were placed in 4% formaldehyde for at least 48 hours before dehydration, paraffin inclusion, and sectioning. Routine hematoxylin eosin saffron (HES) staining was performed and analyzed by a pathologist. Subsequent immunohistological staining of samples were done following clinical protocols by CHU Sainte-Justine’s clinical histology department. For immunofluorescent staining, whole organs or tumor tissues were flash frozen on dry ice after harvest. 6-10 μm-thick sections were made on a cryostat (Leica) and deposited on gelatinized slides and immediately fixed and permeabilized in 95% EtOH. Immunofluorescent staining was performed against human CD45 (Cell signaling, clone D9M8I), human CD8 (Biolegend, clone HIT8a), SV40 LargeT antigen (Santa Cruz, clone Pab 101), and GD2 (Biolegend, clone 14G2a) with AlexaFluor 488 or 594 secondary antibodies and DAPI counterstain.

### Immune infiltrate characterization

Tumors were excised from mice and peripheral blood was harvested at sacrifice. Tumors were digested using the human Tumor Dissociation Kit (with enzymes H and A only to avoid epitope losses) and the GentelMACS Octo Dissociation with heaters (Miltenyi). Cells were then filtered on 70 μm MACS SmartStrainers (Miltenyi) and washed as per manufacturer’s protocol using RPMI (Wisent) with 10% FBS. Cells were then stained with antibodies for analysis by flow cytometry of tumor infiltrating immune cells (hTIICs) against the following targets: humanCD3-AF700, humanCD33-BV510 and humanCD25-BV711 from Biolegend and mouseCD45-PE/Cy7, humanCD45-BUV395, humanCD19-PE/CF594, humanCD4-BB515, humanCD8-BV421, humanCD14-APC/H7, humanCD56-BV786, humanCD127-BB700, humanPD-1-BUV737 and humanTim3-PE from BD Biosciences. Blood samples from the same animals were also stained with the same antibody panel before red blood cell lysis using BD Pharm Lyse lysis buffer (BD Biosciences). All results were acquired using a BD LSRFortessa (BD Biosciences). Data analysis was done on FlowJo V10 (FlowJo, LLC). For tSNE dimensionality reduction analysis, tumor and blood immune cells were subsetted and pooled to obtain a representative sample of 20 000 cells. Dimensionality reduction was done using FlowJo’s tSNE implementation and the FlowSOM algorithm (45) was used to identify clusters in an unbiased manner.

### Statistical analysis

Student’s t-test, one and two-way ANOVA with Šídák’s multiple comparison post-tests were done using GraphPad Prism 8.0. \**p*<0.05, \*\**p*<0.01, *** *p*<0.001, \*\*\*\**p*<0.0001.

## REFERENCES

1. Hay M, Thomas DW, Craighead JL, Economides C, Rosenthal J. Clinical development success rates for investigational drugs. Nature Biotechnology 2014;32(1):40–51 doi 10.1038/nbt.2786.

2. Johnson JI, Decker S, Zaharevitz D, Rubinstein LV, Venditti JM, Schepartz S, et al. Relationships between drug activity in NCI preclinical in vitro and in vivo models and early clinical trials. British journal of cancer 2001;84(10):1424–31 doi 10.1054/bjoc.2001.1796.

3. Dunn GP, Old LJ, Schreiber RD. The three Es of cancer immunoediting. Annual review of immunology 2004;22:329–60 doi 10.1146/annurev.immunol.22.012703.104803.

4. Schreiber RD, Old LJ, Smyth MJ. Cancer immunoediting: integrating immunity’s roles in cancer suppression and promotion. Science (New York, NY) 2011;331(6024):1565–70 doi 10.1126/science.1203486.

5. Galluzzi L, Senovilla L, Zitvogel L, Kroemer G. The secret ally: immunostimulation by anticancer drugs. Nature reviews Drug discovery 2012;11(3):215–33 doi 10.1038/nrd3626.

6. Galluzzi L, Buqué A, Kepp O, Zitvogel L, Kroemer G. Immunological Effects of Conventional Chemotherapy and Targeted Anticancer Agents. Cancer cell 2015;28(6):690–714 doi 10.1016/j.ccell.2015.10.012.

7. Mestas J, Hughes CCW. Of mice and not men: differences between mouse and human immunology. The Journal of Immunology 2004;172(5):2731–8 doi 10.4049/jimmunol.172.5.2731.

8. Balmain A, C.Harris C. Carcinogenesis in mouse and human cells: parallels and paradoxes. Carcinogenesis 2000;21(3):371–7 doi 10.1093/carcin/21.3.371.

9. Hackam DG, Redelmeier DA. Translation of research evidence from animals to humans. JAMA : the journal of the American Medical Association 2006;296(14):1727–32.

10. Fu J, Sen R, Masica DL, Karchin R, Pardoll D, Walter V, et al. Autologous reconstitution of human cancer and immune system in vivo. Oncotarget 2017;8(2):2053–68 doi 10.18632/oncotarget.14026.

11. Jespersen H, Lindberg MF, Nature … D-M. Clinical responses to adoptive T-cell transfer can be modeled in an autologous immune-humanized mouse model. Nature … 2017.

12. Ben-David U, Siranosian B, Ha G, Tang H, Oren Y, Hinohara K, et al. Genetic and transcriptional evolution alters cancer cell line drug response. Nature 2018;560(7718):325–30 doi 10.1038/s41586-018-0409-3.

13. Ben-David U, Beroukhim R, Golub TR. Genomic evolution of cancer models: perils and opportunities. Nature reviews Cancer 2018 doi 10.1038/s41568-018-0095-3.

14. Wang M, Yao LC, Cheng M, Cai D, Martinek J, Pan CX, et al. Humanized mice in studying efficacy and mechanisms of PD-1-targeted cancer immunotherapy. FASEB J 2018;32(3):1537–49 doi 10.1096/fj.201700740R.

15. Zitvogel L, Pitt JM, Daillère R, Smyth MJ, Kroemer G. Mouse models in oncoimmunology. Nature reviews Cancer 2016;16(12):759–73 doi 10.1038/nrc.2016.91.

16. King M, Pearson T, Shultz LD, Leif J, Bottino R, Trucco M, et al. A new Hu-PBL model for the study of human islet alloreactivity based on NOD-scid mice bearing a targeted mutation in the IL-2 receptor gamma chain gene. Clinical immunology 2008;126(3):303–14 doi 10.1016/j.clim.2007.11.001.

17. Marodon G, Desjardins D, Mercey L, Baillou C, Parent P, Manuel M, et al. High diversity of the immune repertoire in humanized NOD.SCID.gamma c-/- mice. European journal of immunology 2009;39(8):2136–45 doi 10.1002/eji.200939480.

18. Kooreman NG, de Almeida PE, Stack JP, Nelakanti RV, Diecke S, Shao N-Y, et al. Alloimmune Responses of Humanized Mice to Human Pluripotent Stem Cell Therapeutics. Cell reports 2017;20(8):1978–90 doi 10.1016/j.celrep.2017.08.003.

19. Denton PW, Krisko JF, Powell DA, Mathias M, Kwak YT, Martinez-Torres F, et al. Systemic administration of antiretrovirals prior to exposure prevents rectal and intravenous HIV-1 transmission in humanized BLT mice. PloS one 2010;5(1) doi 10.1371/journal.pone.0008829.

20. Wahl A, Garcia VJ. The use of BLT humanized mice to investigate the immune reconstitution of the gastrointestinal tract. Journal of Immunological Methods 2014;410:28–33 doi 10.1016/j.jim.2014.06.009.

21. Hahn WC, Counter CM, Lundberg AS, Beijersbergen RL, Brooks MW, Weinberg RA. Creation of human tumour cells with defined genetic elements. Nature 1999;400(6743):464–8 doi 10.1038/22780.

22. Benabdallah B, Desaulniers-Langevin C, Goyer ML, Colas C, Maltais C, Li Y, et al. Myogenic progenitor cells derived from human induced pluripotent stem cell are immune-tolerated in humanized mice. Stem Cells Transl Med 2020 doi 10.1002/sctm.19-0452.

23. Moquin-Beaudry G, Colas C, Li Y, Bazin R, Guimond JV, Haddad E, et al. The Tumor-Immune Response Is Not Compromised by Mesenchymal Stromal Cells in Humanized Mice. J Immunol 2019;203(10):2735–45 doi 10.4049/jimmunol.1900807.

24. Raggi C, M’Callum M-A, Pham QT, Gaub P, Selleri S, Baratang N, et al. Leveraging complex interactions between signaling pathways involved in liver development to robustly improve the maturity and yield of pluripotent stem cell-derived hepatocytes. bioRxiv 2020:2020.09.02.280172 doi 10.1101/2020.09.02.280172.

25. Putra J, Ornvold K. Undifferentiated embryonal sarcoma of the liver: a concise review. Archives of Pathology and Laboratory … 2015.

26. Rich JN, Guo C, McLendon RE, Bigner DD, Wang XF, Counter CM. A genetically tractable model of human glioma formation. Cancer research 2001;61(9):3556–60.

27. Coelho MA, de Carne Trecesson S, Rana S, Zecchin D, Moore C, Molina-Arcas M, et al. Oncogenic RAS Signaling Promotes Tumor Immunoresistance by Stabilizing PD-L1 mRNA. Immunity 2017;47(6):1083–99 e6 doi 10.1016/j.immuni.2017.11.016.

28. Nature V-JE. Cells of origin in cancer. Nature 2011.

29. Singh SK, Hawkins C, Clarke ID, Squire JA, Bayani J, Hide T, et al. Identification of human brain tumour initiating cells. Nature 2004;432(7015):396–401 doi 10.1038/nature03128.

30. Bailey MH, Tokheim C, Porta-Pardo E, Cell S-S. Comprehensive characterization of cancer driver genes and mutations. Cell 2018.

31. Smith RC, Tabar V. Constructing and Deconstructing Cancers using Human Pluripotent Stem Cells and Organoids. Cell Stem Cell 2019;24(1):12–24 doi 10.1016/j.stem.2018.11.012.

32. Koga T, Chaim IA, Benitez JA, Markmiller S, Parisian AD, Hevner RF, et al. Longitudinal assessment of tumor development using cancer avatars derived from genetically engineered pluripotent stem cells. Nat Commun 2020;11(1):550 doi 10.1038/s41467-020-14312-1.

33. Brehm MA, Kenney LL, Wiles MV, Low BE, Tisch RM, Burzenski L, et al. Lack of acute xenogeneic graft-versus-host disease, but retention of T-cell function following engraftment of human peripheral blood mononuclear cells in NSG mice deficient in MHC class I and II expression. FASEB J 2018:fj201800636R doi 10.1096/fj.201800636R.

34. Ashizawa T, Iizuka A, Nonomura C, Kondou R, Maeda C, Miyata H, et al. Antitumor Effect of Programmed Death-1 (PD-1) Blockade in Humanized the NOG-MHC Double Knockout Mouse. Clin Cancer Res 2017;23(1):149–58 doi 10.1158/1078-0432.CCR-16-0122.

35. Tatum AM, Mylin LM, Bender SJ, Fischer MA, Vigliotti BA, Tevethia MJ, et al. CD8+ T cells targeting a single immunodominant epitope are sufficient for elimination of established SV40 T antigen-induced brain tumors. Journal of immunology (Baltimore, Md : 1950) 2008;181(6):4406–17 doi 10.4049/jimmunol.181.6.4406.

36. Kooreman NG, Kim Y, de Almeida PE, Termglinchan V, Diecke S, Shao NY, et al. Autologous iPSC-Based Vaccines Elicit Anti-tumor Responses In Vivo. Cell Stem Cell 2018;22(4):501–13 e7 doi 10.1016/j.stem.2018.01.016.

37. Huntington ND, Legrand N, Alves NL, Jaron B, Weijer K, Plet A, et al. IL-15 trans-presentation promotes human NK cell development and differentiation in vivo. The Journal of experimental medicine 2009;206(1):25–34 doi 10.1084/jem.20082013.

38. Herndler-Brandstetter D, Shan L, Yao Y, Stecher C, Plajer V, Lietzenmayer M, et al. Humanized mouse model supports development, function, and tissue residency of human natural killer cells. Proceedings of the National Academy of Sciences of the United States of America 2017;114(45):E9626–E34 doi 10.1073/pnas.1705301114.

39. Katano I, Takahashi T, Ito R, Kamisako T, Mizusawa T, Ka Y, et al. Predominant Development of Mature and Functional Human NK Cells in a Novel Human IL-2–Producing Transgenic NOG Mouse. The Journal of Immunology 2015;194(7):3513–25 doi 10.4049/jimmunol.1401323.

40. Wege AK, Melkus MW, Denton PW, Estes JD, Garcia JV. Functional and phenotypic characterization of the humanized BLT mouse model. Current topics in microbiology and immunology 2008;324:149–65 doi 10.1007/978-3-540-75647-7_10.

41. Benabdallah B, Désaulniers-Langevin C, Colas C, Li Y, Rousseau G, Guimond JV, et al. Natural Killer Cells Prevent the Formation of Teratomas Derived From Human Induced Pluripotent Stem Cells. Frontiers in immunology 2019;10(2580) doi 10.3389/fimmu.2019.02580.

42. Beauséjour CM, Krtolica A, Galimi F, Narita M, Lowe SW, Yaswen P, et al. Reversal of human cellular senescence: roles of the p53 and p16 pathways. The EMBO journal 2003;22(16):4212–22 doi 10.1093/emboj/cdg417.

43. Beier KT, Samson ME, Matsuda T, Cepko CL. Conditional expression of the TVA receptor allows clonal analysis of descendents from Cre-expressing progenitor cells. Developmental biology 2011;353(2):309–20 doi 10.1016/j.ydbio.2011.03.004.

44. Shultz LD, Brehm MA, Garcia-Martinez JV, Greiner DL. Humanized mice for immune system investigation: progress, promise and challenges. Nat Rev Immunol 2012;12(11):786–98 doi 10.1038/nri3311.

45. Gassen S, Callebaut B, Helden MJ, Lambrecht BN, Demeester P, Dhaene T, et al. FlowSOM: Using self-organizing maps for visualization and interpretation of cytometry data. Cytometry Part A 2015;87(7):636–45 doi 10.1002/cyto.a.22625.

