## Supplementary material for "Autologous humanized mouse models of iPSC-derived tumors allow for the evaluation and modulation of cancer-immune cell interactions": supp figure 1-9

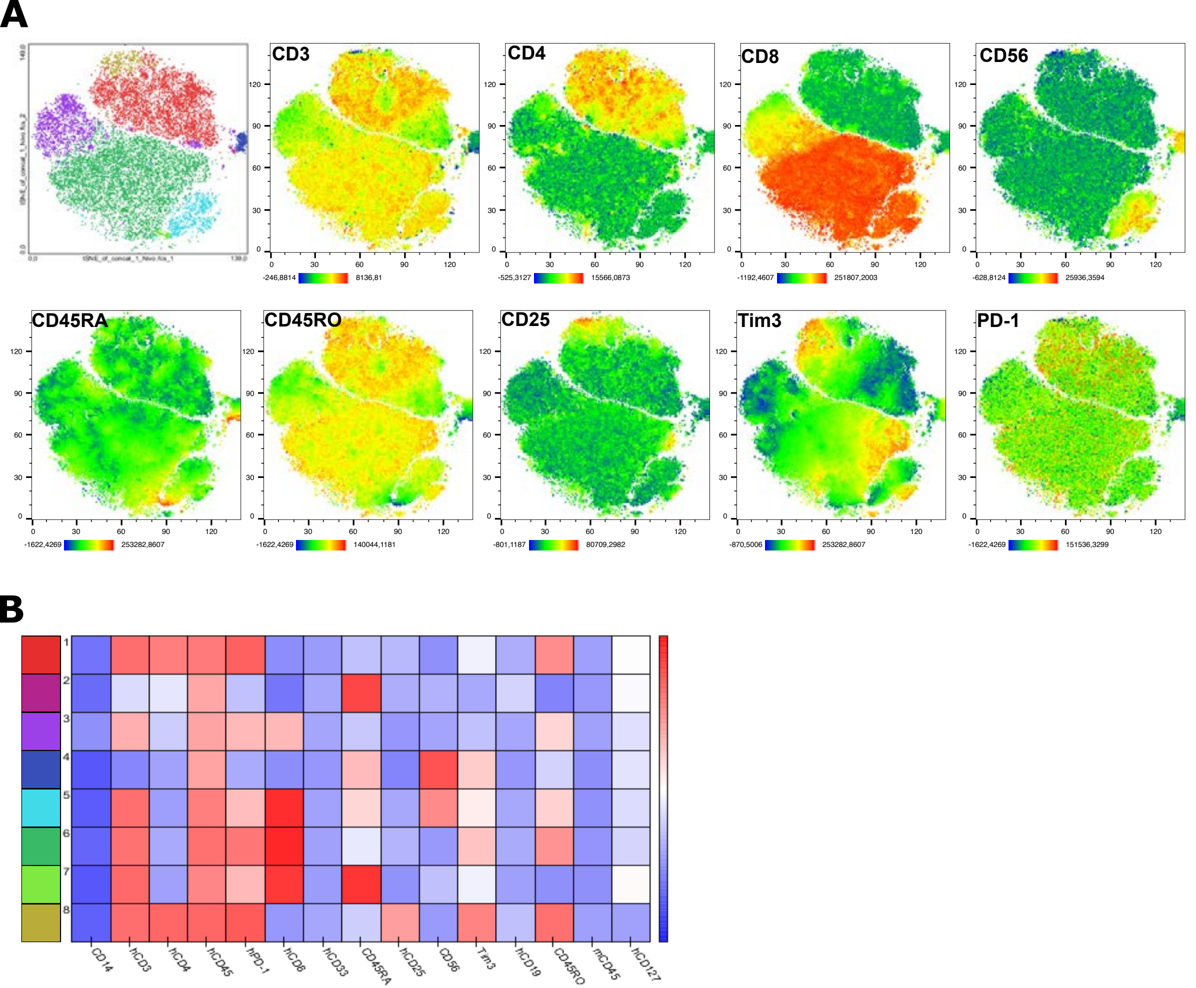


**Figure S1. Human immune population clustering and protein expression in Auto-AT mice blood and tumors.** (A) tSNE dimensional reduction plots of human immune compartment of Auto-AT mice blood and tumors showing unbiased cluster identification (top leftmost) and flow cytometry protein detection heatmaps. (B) Cluster protein expression heatmap for reference clusters in (A) top leftmost panel.

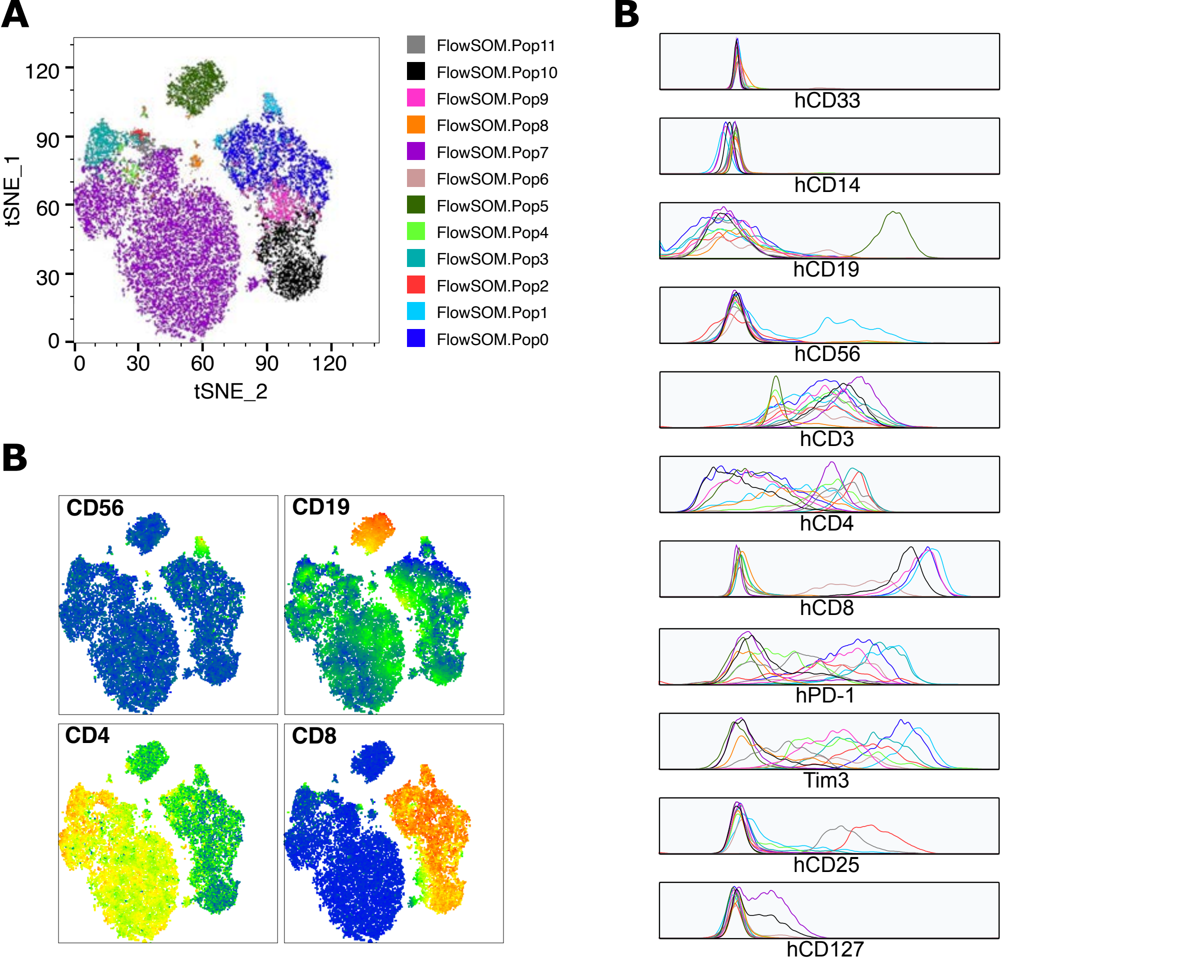


**Figure S2. Auto-BLT human immune cell clustering.** (A) tSNE dimentional reduction plot with cluster annotation as obtained by FlowSOM unsupervised clustering tool in FlowJo. (B) Detailed fluorescence histograms for differential protein expression of individual clusters in pooled cells collected from blood and tumor. (C) Gene expression heatmap for principal cell population clusters.

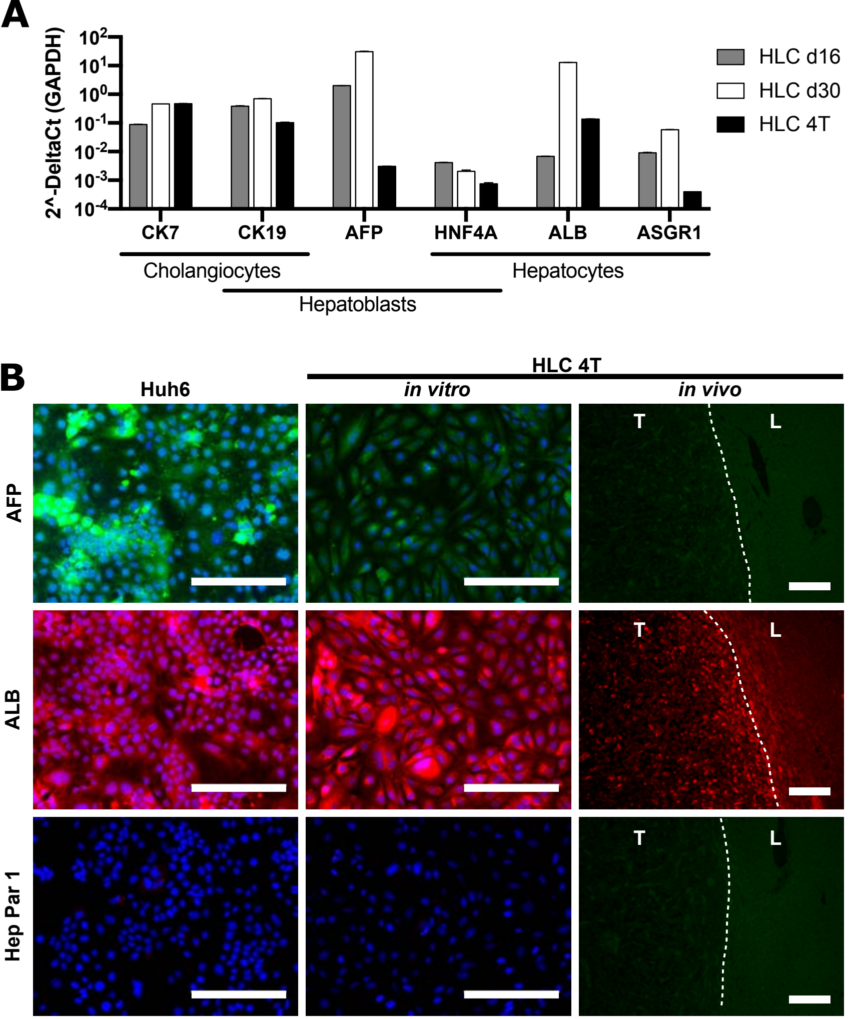


**Figure S3. Characterization of hepatic differentiation markers in HLC cells.** (A) RT-qPCR gene expression analysis for untransformed HLC at days 16 and 30 of differentiation and 4T transformed HLC (passage 2). CK7 cytokeratin 7, CK19; cytokeratin 19, AFP; Alpha fetoprotein, HNF4A; Hepatic nuclear factor 4 alpha, ALB; albumin, ASGR1; asialoglycoprotein receptor 1. (B) Immunofluorescent staining for AFP, ALB and Hepatocyte paraffin 1 antibody (Hep Par 1) of HuH6 hepatoblastoma cell line in vitro and HLC 4T cell line in vitro and in vivo. Huh6 are positive for AFP and ALB but negative for Hep Par 1. In vitro, HLC 4T are weakly positive for AFP, negative for Hep Par1 and positive for ALB. In vivo, HLC 4T are negative for AFP and Hep Par 1 and weakly positive for ALB. Tumor (T) and liver parenchyma (L) separated by dotted line (right). All scale bars = 200 µm.

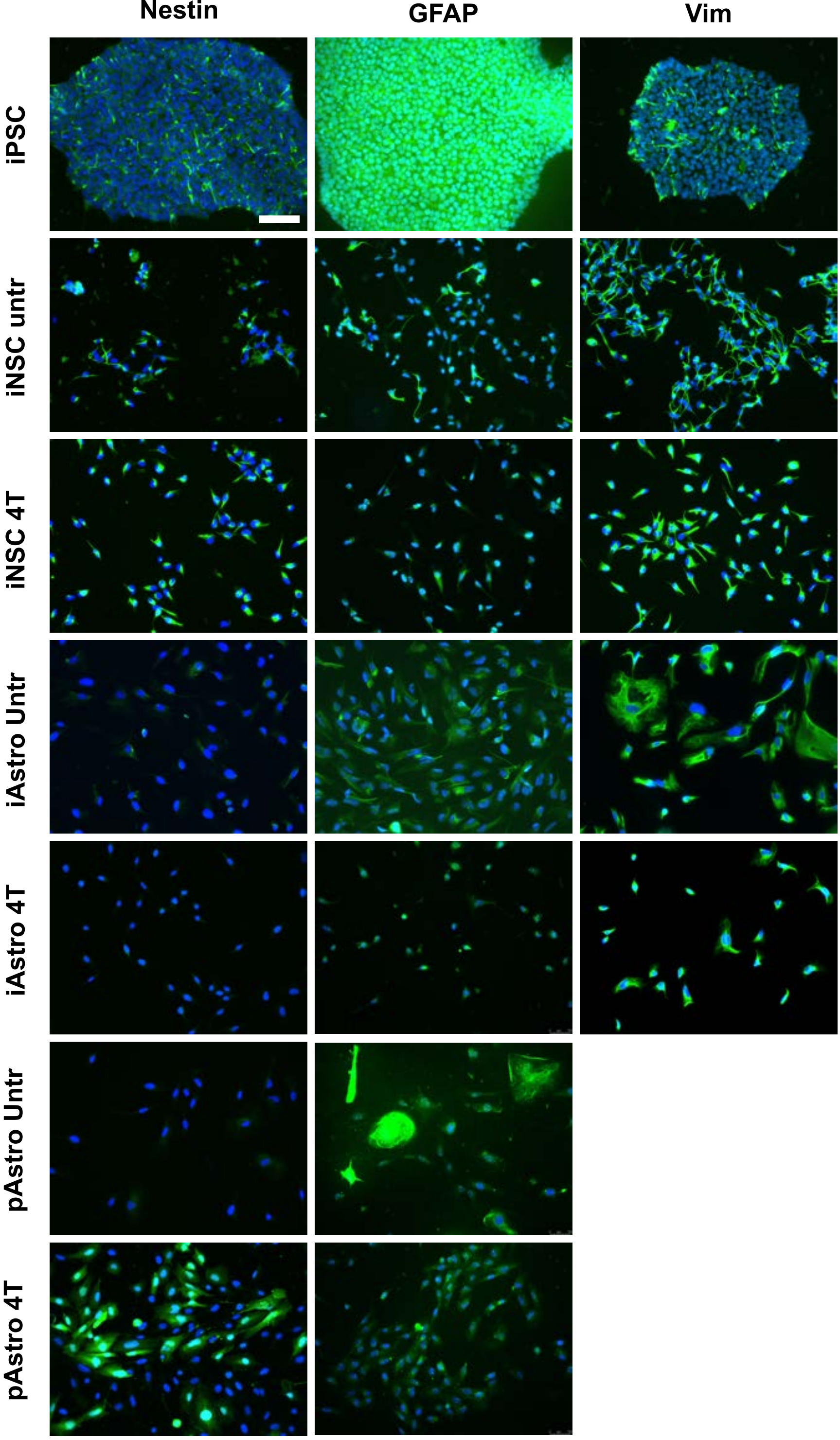


**Figure S4. Characterization of differentiation markers in iNSC and iAstro cell lines.** Expression of nestin, glial fibrillary acidic protein (GFAP) and vimentin in multiple cell populations. Identical acquisition parameters were employed for each staining in all samples. As expected, NSC but not astrocytic populations expressed nestin with the exception of transformed primary astrocytes (pAstro 4T). Of note, such a high GFAP expression in the iPSC population was unexpected. The detection of some GFAP positive cells in iNSC and iNSC 4T populations may be attributable to spontaneous differentiation, or lack thereof given parental iPSC were positive for GFAP. We also observed a general decrease of GFAP expression in transformed iNSC 4T cell lines. Vimentin is strongly expressed by all populations upon differentiation and is maintained after transformation. iNSC = iPSC-derived NSC, iAstro = iPSC-derived astrocytes, pAstro = primary astrocytes. Scale bar = 200μm

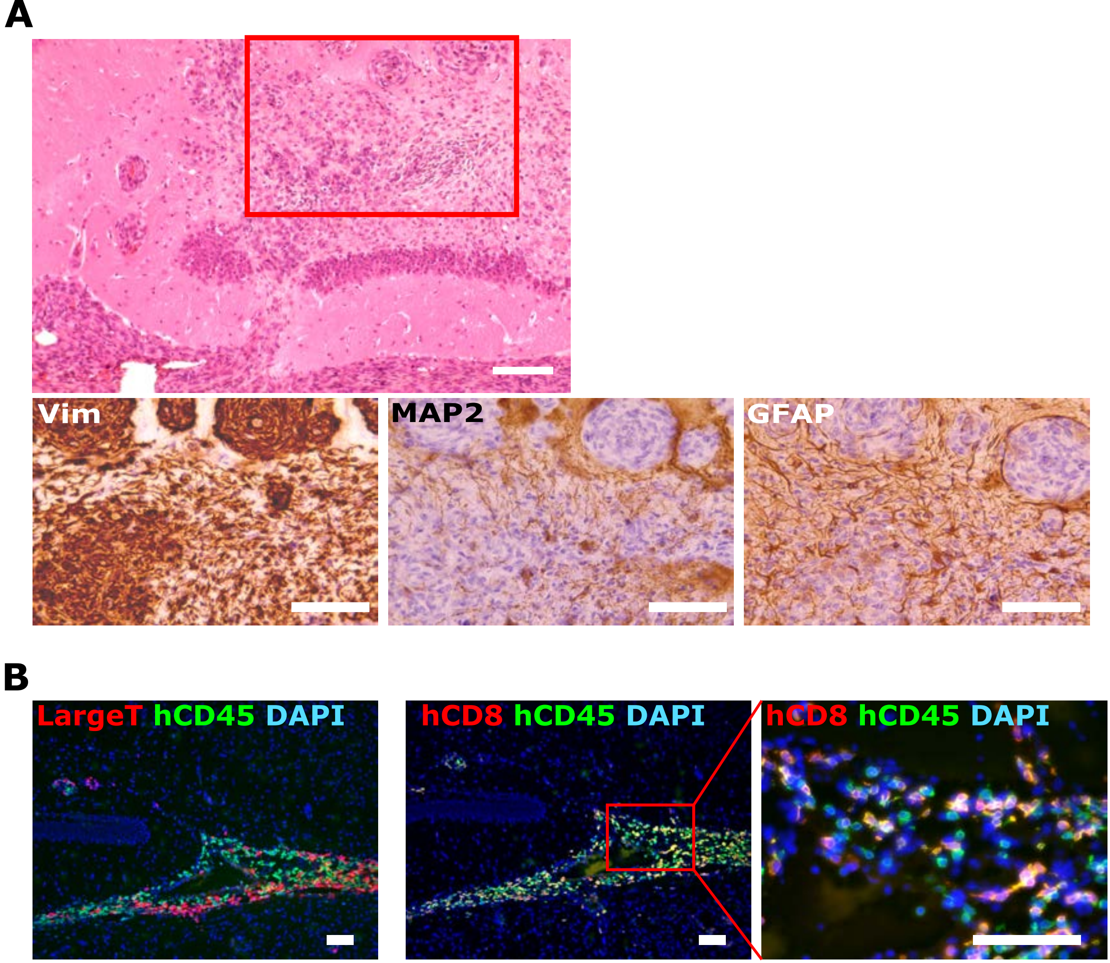


**Figure S5. Phenotypic characterization of iNSC 4T tumors.** (A) Representative photomicrograph of an iNSC 4T tumor isolated on day 21 after orthotopic injection. High magnification images show tumor cells growing near the dentate gyrus (top) and expressing Vimentin, but low to no expression of MAP2 and GFAP (bottom). Most GFAP positive staining is attributed to reactive mouse astrocytes. red rectangle = high magnification region. (B) Immunofluorescent staining of tumor immune cell infiltration showing specific accumulation of hCD45 (green) cells within tumor masses (left, Large T antigen in red) with a high proportion of CD8 positive T cells (middle and right, red). Scale bars = 100µm

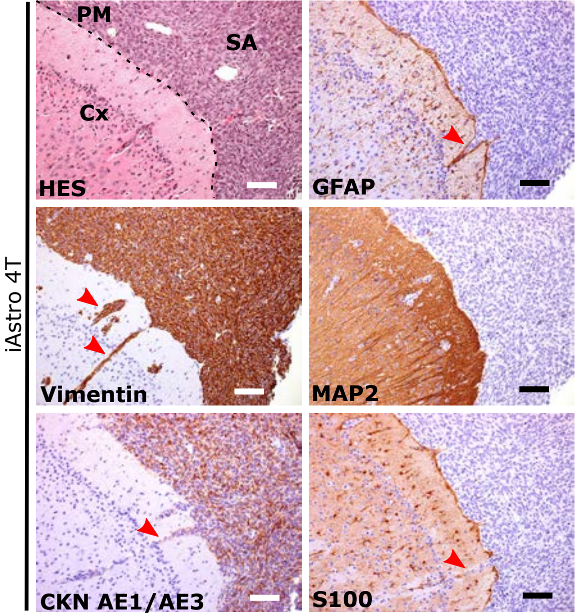


**Figure S6. iAstro 4T tumor molecular characterization.** Subarachnoid focus of a representative iAstro 4T tumor stained with hematoxylin eosin saffron (HES), vimentin, cytokeratin AE1/AE3, glial fibrillary acidic protein (GFAP), microtubule-associated protein 2 (MAP2) and S100 protein. Note the strong tumor cell immunoreactivity for vimentin and cytokeratin and negative staining for GFAP, MAP2 and S100. Dotted line represents pia mater (PM) with cortex on the left (Cx) and tumor-invaded subarachnoid space (SA), and red arrowheads indicate tumor-infiltrated Virchow-Robin spaces.

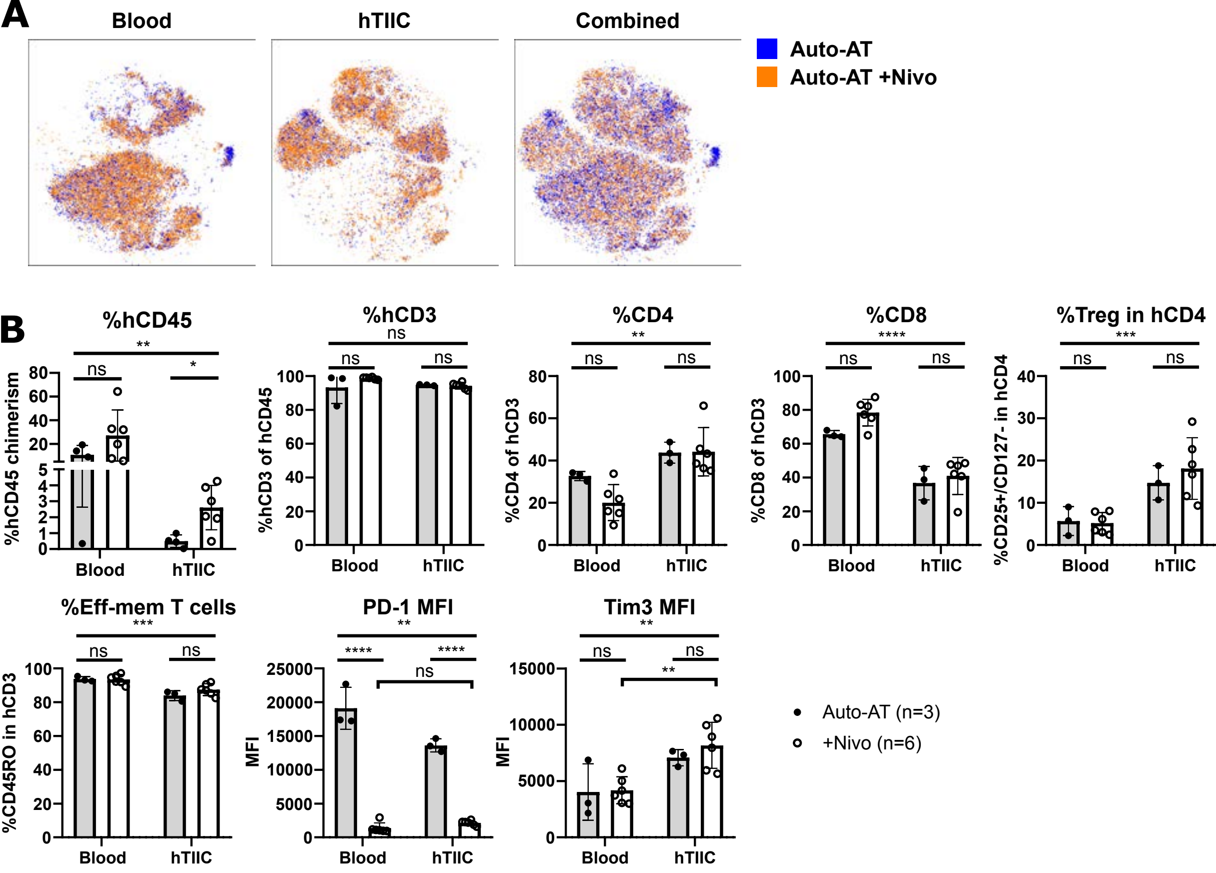


**Figure S7. Nivolumab impact on immune cell phenotype in blood and fibroblastic tumors.** (A) tSNE dimensional reduction of human immune cells by flow cytometry. Blood (left), tumor infiltrating (middle) and combined (right) human immune cells in Auto-AT mice with (orange) or without (blue). The Nivolumab injected group shows nearly complete overlap of all populations, indicating no significant shift is observed. (B) Flow cytometric quantification of human immune population frequencies (top row) and T cell phenotype (bottom row) in blood and tumors of Auto-AT alone (grey bars and full dots) or with Nivolumab treatment (hollow bars and dots). Mean±SD. One tumor per mouse was processed. Note, the almost complete absence of PD-1 expression in both populations is likely the result of the interference of Nivolumab in the staining.

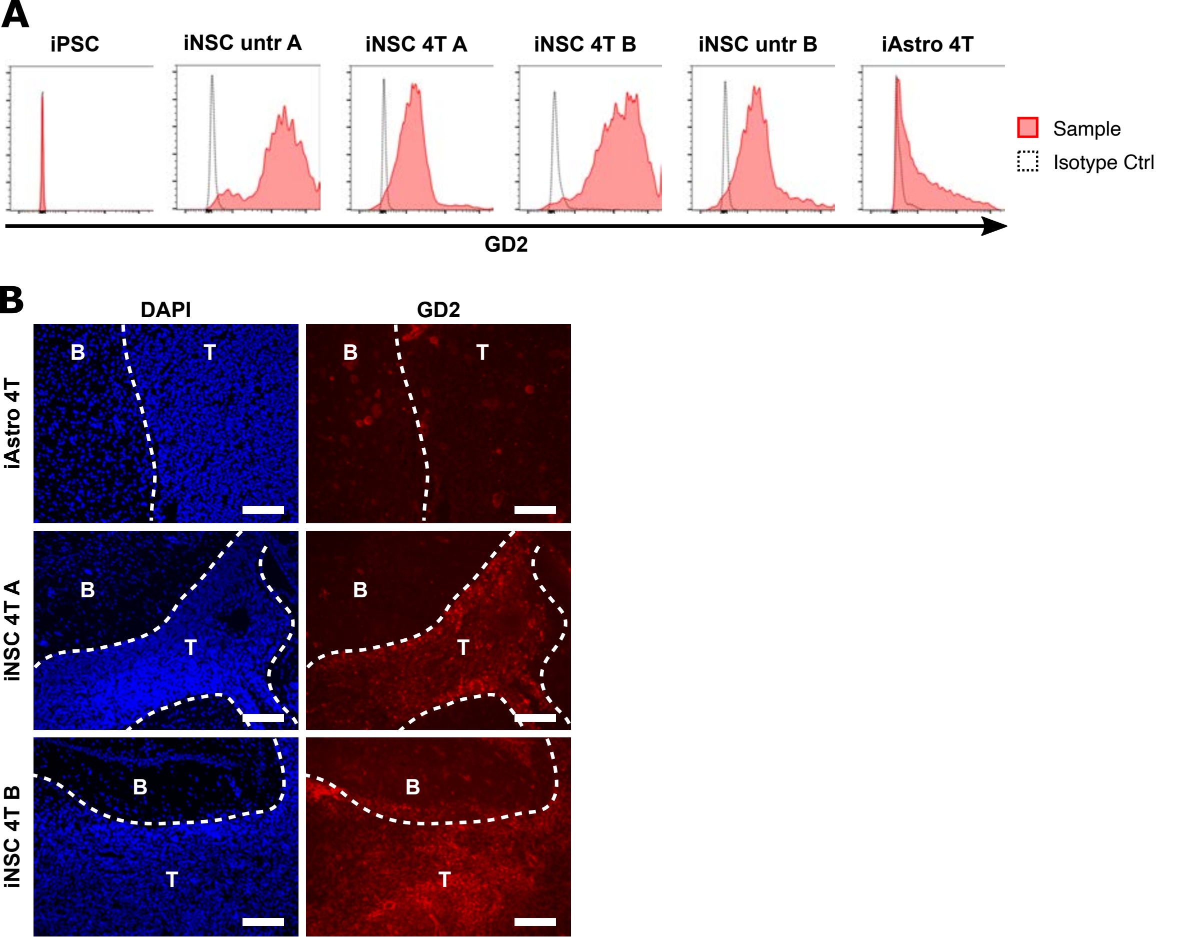


**Figure S8. Expression of the tumor specific antigen GD2 by iNSC tumor cells** (A) Flow cytometric assessment of GD2 expression in different steps of iPSC differentiation and transformation showing specific expression of GD2 in normal and transformed iNSCs for donors A and B. (B) Immunofluorescent staining of in vivo iAstro (top) and iNSC tumors showing little or no expression of GD2 in iAstro 4T tumor (top) and specific expression of GD2 in iNSC tumors (T) from both donors A (center) and B (bottom). Blue = DAPI counterstain, red = GD2, dotted line = tumor edge, B = brain parenchyma, T = tumor tissue.

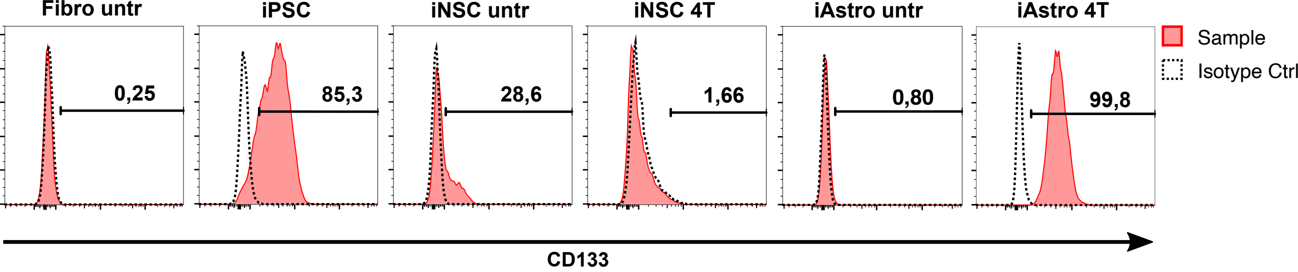


**Figure S9. CD133 expression in iPSC-derived astrocytic lineages.** Detection of the glioblastoma cancer stem cell marker CD133 by flow cytometry in iAstro 4T and iPSC. No significant expression was detected in primary fibroblasts from the original donor (Fibro untr), transformed iNSC or untransformed iAstro. Of note, a low-expressing subpopulation was observed in untransformed iNSC. Red = anti-CD133 staining, dotted line = isotype control.
